# Allergen-responsive T helper type 2 cells revealed by high-dimensional profiling in allergen challenged human airways

**DOI:** 10.64898/2026.07.31.741782

**Authors:** Benjamin D. Wheeler, Jingming Wang, Srilaxmi Nerella, Kristina Johansson, Suresh Garudadri, Priscila Muñoz-Sandoval, Thomas Mazumder, Stephanie A. Christenson, Nirav R. Bhakta, David J. Erle, K. Mark Ansel, Prescott G. Woodruff

## Abstract

Asthma is a chronic inflammatory disease affecting over 300 million people worldwide. This disease has multiple underlying etiologies, and a major endotype of asthma is characterized by cellular and molecular signatures of type 2 (allergic) inflammation. In this study we conducted bronchoscopies with airway segmental allergen challenge in allergic asthmatics to dissect airway responses to allergen. Using mass cytometry and single-cell RNA sequencing, we characterized with high resolution the airway immune landscape before and after allergen challenge and the heterogeneity present between subjects. This heterogeneity generally falls along a type 1/ type 2 axis. In type 2 high individuals, we identified allergen-reactive Th2 cells by using TCR sequences to barcode clonal T cell populations in single-cell genomic and activation-induced marker expression assays. These potentially pathogenic Th2 cell clones were present systemically and expanded following allergen challenge, connecting local lung inflammation to systemic clonal Th2 cell dynamics. Th2 cell airway ingress was coordinated with myeloid cell expression of T cell chemoattractants including CCL17 and CCL22. This study provides insight into the molecular and cellular components of allergen-induced tissue inflammation in asthma. Deeper resolution of the T cell response to aeroallergens may inform novel diagnostic and therapeutic strategies for asthma and other allergic airway diseases.

## Introduction

Asthma is a chronic disease of the airways affecting 334 million people with 250,000 annual deaths worldwide (GBD Chronic Respiratory Disease Collaborators 2020). Asthma is characterized by inflammation in the airways leading to reversible airway obstruction that can cause coughing, wheezing, chest tightness, and other symptoms (Padem & Saltoun 2019). In addition to episodic bronchospasm and heightened inflammation, other hallmark features of asthma include excess mucus production and airway remodeling that can lead to chronic obstruction and impaired lung function (Tang et al. 2022). To address these issues, patients are typically treated with corticosteroids (inhaled or systemic) and bronchodilators (Lin et al. 2018). The therapeutic landscape for these patients has broadened in recent years with the advent of biologics directly targeting cytokines involved in allergic (type 2) inflammation, including TSLP, IL-5, and IL-4 and IL-13 via their shared receptor IL-4RA (Wenzel et al. 2016; Gauvreau et al. 2014; Menzies-Gow et al. 2003). However, these therapies are expensive, must be continued indefinitely, and are only effective in a subset of individuals with asthma (Moran & Pavord 2020; Hansen et al. 2024) .

The success of these therapies reinforces the importance of type 2 inflammation as a critical component of asthma pathology. However, there is significant heterogeneity present in this patient population (Fitzpatrick et al. 2011; Haldar et al. 2008; Moore et al. 2010). The type 2 high asthma endotype is characterized by increased expression of the IL-13 responsive genes *POSTN*, *CLCA1*, and *SERPINB2* in lung epithelium (Bhakta et al. 2013; Woodruff et al. 2009). Type 2 high asthma is also typically marked by elevated blood and sputum eosinophils (Dunican et al. 2018; Castro et al. 2018) and increased serum IgE, both of which contribute to asthma pathology and are driven by type 2 cytokines. However, the identification of type 2 cytokine secreting T helper cells in the lung has been difficult.

It is important to further understand how allergen exposure induces allergic responses in the local lung environment. To address this gap, we performed high dimensional mass cytometry and transcriptomic analysis of airway cells in a clinical study of segmental airway allergen challenge. These analyses revealed that type 2 helper T cells are specifically allergen responsive and elevated in subjects with other characteristics of type 2 inflammation including airway eosinophilia, allergen-specific blood IgE, and elevated expression of type 2 responsive genes in airway epithelial cells (AECs). Integrative analysis of these data sets revealed inflammatory axes present between infiltrating monocyte-derived populations, the chemokine environment they establish, and the resultant ingress of T cells and other lymphocyte populations.

## Results

### High dimensional cytometry defines the local type 2 inflammatory response to allergen challenge

We probed the local response to allergen exposure in a study of segmental allergen challenge. The study was conducted over the course of three clinical visits, schematized in [Figure 1a]. Subjects were enrolled with stable or well-controlled asthma, not on inhaled corticosteroids. Enrolled subjects with asthma were between the ages of 18 to 50, had a baseline FEV1 > 75% of predicted, a methacholine PC_20_ < 16 mg/mL, and skin reactivity to either house dust mit*e Dermatophagoides pteronyssinus* (HDM) or cat dander (Table 1). In the first clinical visit (V1), to assess allergen specific responses, subjects first underwent qualitative skin prick testing to determine whether the participant responds to HDM or cat dander (cat). Then, in the same V1, quantitative skin prick testing was done with the allergen to which they responded to determine the threshold allergen concentration which elicits an allergic response. In the second visit (V2), the following experimental baseline samples were collected: peripheral blood for whole blood RNA, spirometry, and via fiberoptic bronchoscopy “baseline” (BL) epithelial brushing in the left lower lobe and bronchoalveolar lavage (BAL) in left upper lobe. Then, a dose of allergen calibrated to the threshold dose in skin prick testing (the full threshold skin dose for cat and one third of the threshold skin dose for HDM) was instilled into the a segment of either the right middle lobe (RML) or right upper lobe (RUL, selected randomly) and PBS diluent was administered in either the RML or RUL (the lobe which did not receive allergen). The third visit (V3) was performed either 1 day (early) or 7 days (late) after V2 and BAL and epithelial brushings were performed from the sites of allergen and diluent challenge, yielding paired allergen challenged (AC) and diluent challenged (DIL) samples from each subject.

Consistent with prior reports, allergen challenge induced robust airway inflammation [Figure 1b]. The gross cellular composition of BAL cells was significantly changed by allergen challenge [Figure 1c,d]. At both 1d and 7d after challenge, this change was dominated by an increase in eosinophils that resulted in a proportional decrease in macrophages, the dominant population in BL samples. While eosinophil recruitment was a prominent response to allergen, we observed significant heterogeneity between subjects, with some experiencing neutrophil and/or lymphocyte infiltration. There was no clear relationship between eosinophil and neutrophil proportions in the BL and DIL samples [Figure 1e]. However, there was a strong inverse correlation between the proportion of BAL eosinophils and neutrophils in AC samples, indicating a polarized response to allergen [Figure 1e].

To deeply characterize the airway-infiltrating immune cell populations, we employed high dimensional cytometry by time of flight (CyTOF) (Nassar et al. 2016). We developed a panel of 34 cell surface markers and used it to distinguish 13 distinct cell types by bivariate gating [Supplemental Figure 1]. Principal component analysis (PCA) of the V3 CyTOF and cytospin data showed a consistent cellular composition in DIL samples, whereas AC samples diverged along two distinct axes [Figure 2a]. Analyzing AC samples alone better resolved this dichotomy, which skewed mainly along PC1, representing 22% of the variance in these samples [Figure 2b].

Examination of the measured variables that contributed to PC1 revealed that the dichotomy between neutrophils and eosinophils observed in the cytospin slides extended to other cell types, indicating modules of type 1 and type 2 inflammation induced by allergen challenge [Figure 2c]. Low PC1 values were driven by high proportions of neutrophils, monocytes, and macrophages [Figure 2c]. AC samples with these characteristics included all 3 AC samples from non-allergic subjects, yet differed from DIL samples from the same subjects, suggesting a significant contribution from the innate immune response to components of the allergen extracts. In contrast, samples with high PC1 were dominated by allergic subjects, including several of the late (day 7) samples [Figure 2b]. These samples contained high eosinophils and elevated proportions of basophils, ILC2s, and B and T lymphocytes [Figure 2c], indicative of an adaptive type 2 response to the allergen challenge. Baseline HDM-specific plasma IgE positively correlated with PC1 in subjects analyzed early (day 1) after HDM challenge [Figure 2d]. Thus, allergen challenge induced local type 2 inflammation especially in subjects with high allergen-specific IgE, a marker of pre-existing systemic type 2 immunity [Figure 2 d].

**Table 1.** Subject Table.

|  |  | Non-Allergic Non-Asthmatics (n = 3) | Allergic Non-Asthmatic (n = 2) | Allergic Asthmatic (n = 19) |
| --- | --- | --- | --- | --- |
| Age, mean [SD] |  | 29.0 [5.0] | 29.5 [2.1] | 29.2 [7.8] |
| Male, n (%) |  | 2 (66%) | 1 (50%) | 6 (32%) |
| Non-white |  | 2 (66%) | 0 (0%) | 11 (58%) |
| Hispanic |  | 0 (0%) | 1 (50%) | 4 (21%) |
| D. Pter Reactive | N (%) | 0 (0%) | 2 (100%) | 19 (100%) |
|  | Skin Reactivity, mm [SD] | 0 [0] | 6.75 [4.2] | 11.1 [5.3] |
| Cat Reactive | N (%) | 0 (0%) | 0 (0%) | 6 (32%) |
|  | Skin Reactivity if reactive, mm [SD] | NA | NA | 7.2 [3.9] |

**Figure 1.**
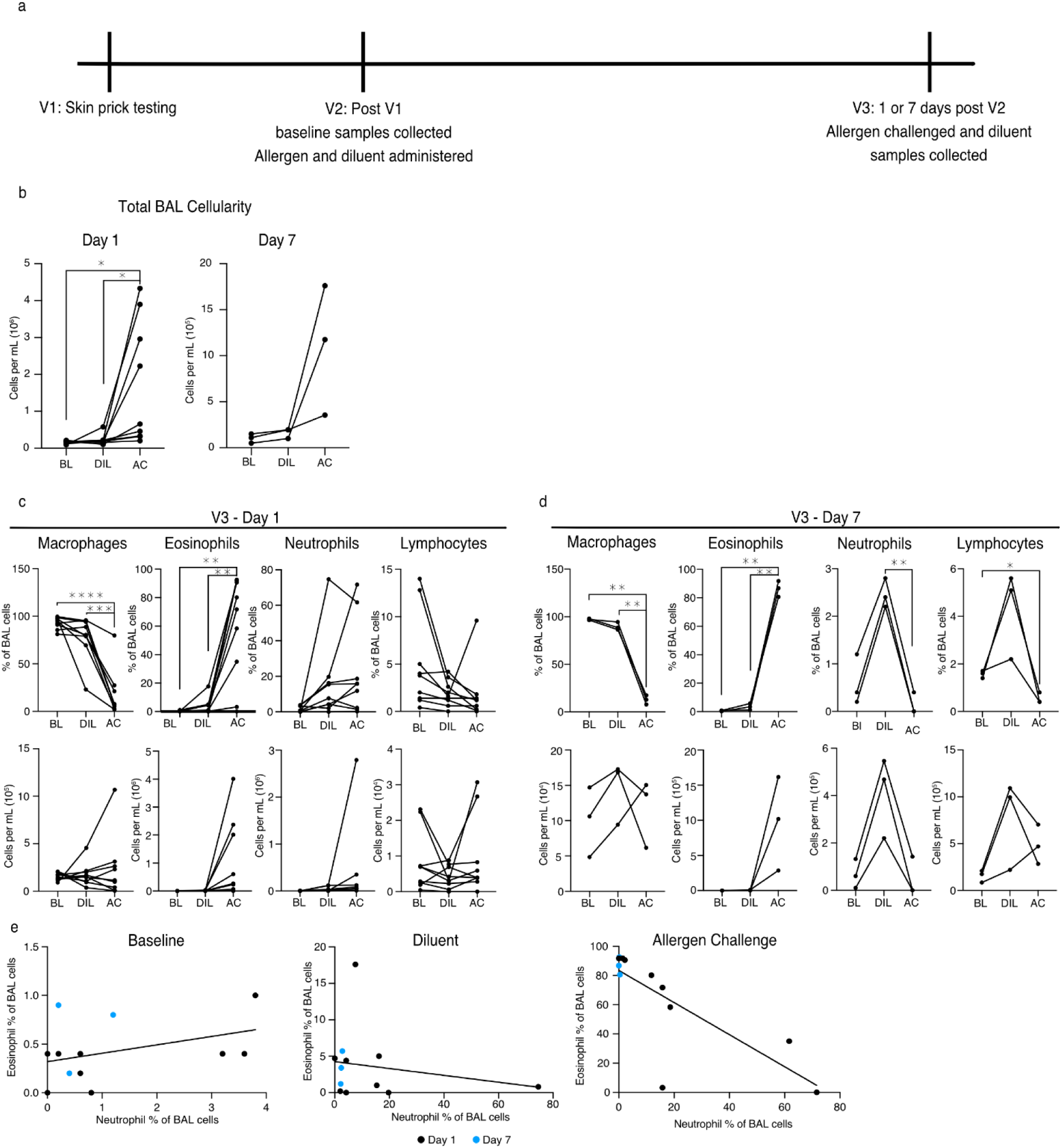
Allergen challenge induces leukocyte infiltration into the airway lumen a. Schematic of study design and clinical visits b. Total BAL cellularity measured by cells per mL of BAL fluid returned from bronchoscopies at all time points for subjects challenged with HDM c. BAL cell composition quantified by cytospin slides from subjects challenged with HDM and part of the day 1 arm of the study as both percent of cells counted and concentration per mL of returned BAL fluid d. BAL cell composition quantified by cytospin slides from subjects challenged with HDM and part of the day 1 arm of the study as both percent of cells counted and concentration per mL of returned BAL fluid e. Comparison of eosinophil and neutrophil composition of BAL fluid by percent of cells counted in cytospin data at the baseline visit, the diluent challenged sample, and the allergen challenged sample for both the day 1 and day 7 samples (BL R^2^: 0.15, p: 0.22; DIL R^2^: 0.04, p: 0.54; AC R^2^: 0.62, p: 0.002). Statistics displayed determined by Dunnett’s multiple comparisons test following one-way anova between AC versus BL and AC versus DIL (*,adjusted p<0.05;**, adjusted p<0.01;***,adjusted p<0.001;****,adjusted p<0.0001)

**Figure 2.**
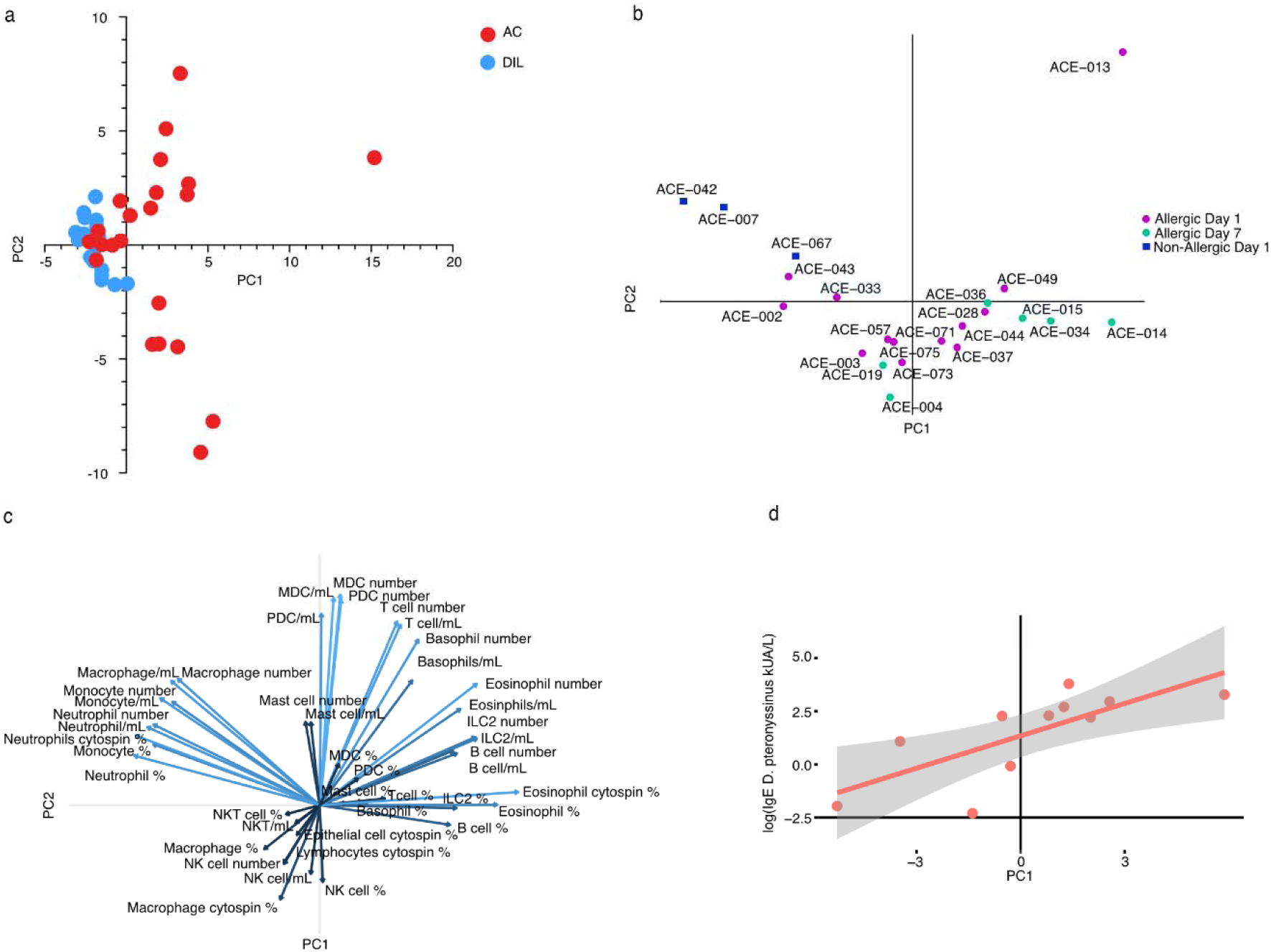
CyTOF resolves Type 1 - Type 2 axis of response to allergen challenge a. Principal component 1 versus principal component 2 of all BAL populations measured by CyTOF and cytospin slides for all individuals, time points, and treatments including those challenged with either cat dander or HDM b. Principal component 1 versus principal component 2 from subsequent analysis of all BAL populations measured by CyTOF and cytospin slides for only the V3 allergen challenged samples including those challenged with either cat dander or HDM c. Vectorized display of how each variable contributes to principal components 1 and 2. Vector shade indicates squared cosine value with lighter shades indicating a higher value subsequently stronger contribution to given principle components. d. Blood concentration of baseline *Dermatophagoides pteronyssinus* specific IgE versus the PC1 value from only HDM allergen challenge for day 1 subjects

### Allergen challenge induces type 2 cytokine response genes in the airway epithelium

The airway epithelium constitutes both a critical barrier surface and an early and active component of the response to allergen exposure (Duchesne et al. 2022; Lambrecht et al. 2019). Using bulk RNA sequencing, we compared gene expression in epithelial brushings collected at baseline and after allergen challenge. To assess the role of cytokine responses, we used gene set enrichment analysis (GSEA) to determine if cytokine-induced gene sets derived from cytokine treatment of air-liquid-interface (ALI) cultures of human bronchial epithelial cells (HBECs) were differentially expressed in AC and BL samples [Figure 3 a-d] (Koh et al. 2023). IL-13 responsive genes were enriched among genes induced by allergen challenge in both allergic asthmatic (AA) and allergic non-asthmatic (ANA) subjects [Figure 3 a&b]. Interestingly, allergen challenge also induced IL-17 responsive gene expression, but only in ANA subjects [Figure 3 c&d].

Given that the IL-17 gene set was only responsive in ANA subjects we sought to more broadly compare allergen-induced gene expression patterns between AA and ANA subjects. Among differentially expressed genes (DEGs) (raw p<0.05) detected between AC and BL brushings in either population, very few were shared between the populations (only 19 out of the total 1026) [Figure 3e]. This trend held when considering DEGs between AC and DIL brushings [Figure 3 e&f]. The DEGs common to both populations revealed no clear functional modules by gene ontology (GO) term analysis [Figure 3g]. However, we did note that IL13RA1 was specifically upregulated by allergen challenge in the AA group, while IL17RA was specifically upregulated in ANA subjects [Figure 3e]

This apparent difference in cytokine response led us to investigate what role individual heterogeneity played in the expression of specific type 2 associated genes in the lung epithelium. Using our RNAseq data, we calculated the 3 gene mean expression of *POSTN*, *CLCA1*, and *SERPINB2*, a measure of type 2 cytokine responses in AECs (Bhakta et al. 2013). This measure was highest among individuals with the highest allergen response-associated CyTOF PC1 score [Figure 3 h-j]. Interestingly, the 3 gene mean was not significantly further induced in AC brushings compared with BL or DIL. Together, these data indicate that AEC gene expression is a stable indicator of type 2 inflammation at baseline that correlates with the magnitude of inflammatory responses to acute allergen challenge. We then hypothesized that this elevated type 2 status would drive additional persistent AEC gene expression changes in these individuals. We subsetted subjects by type 2 high (PC1>0) and type 2 low (PC1<0) status and performed GO term analysis on DEGs detected between these two groups. In BL samples, DEGs elevated in the type 2 high group reflected multiple pathways relating to protein targeting and trafficking, possibly reflective of increased secretory cell composition of the epithelium [Figure 3k]. In AC samples, the most enriched GO terms contained some similar terms (e.g. SRP-dependent cotranslational protein targeting to membrane), but largely consisted of terms related to leukocyte activation and degranulation [Figure 3l]. The enrichment of these terms likely reflects the local infiltration of eosinophils, neutrophils, T cells, and other leukocytes (Alladina et al. 2023).

**Figure 3.**
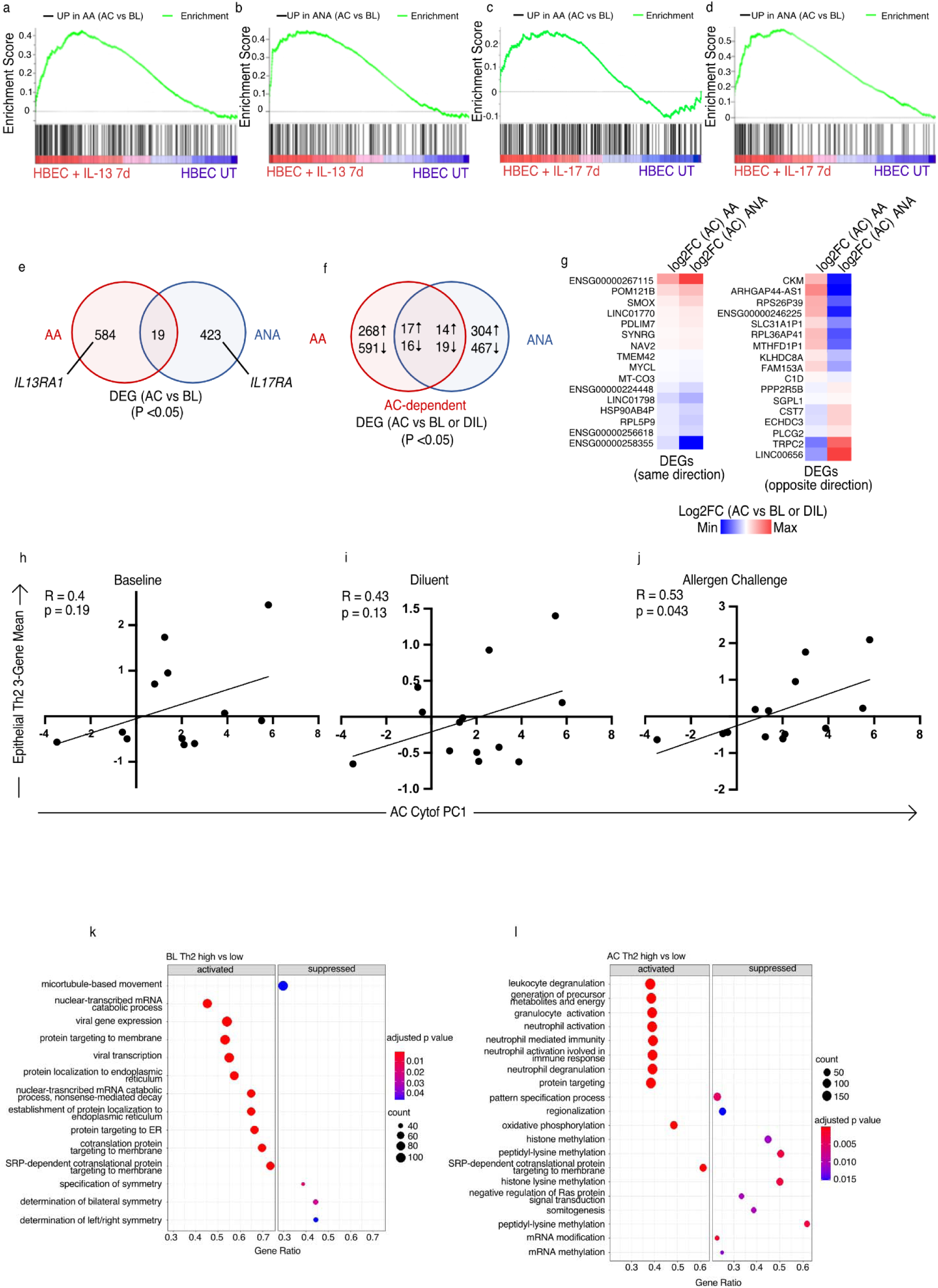
IL-13 responsive genes are induced by allergen challenge and highest in subjects with type 2 high BAL infiltration a. Gene set enrichment analysis (GSEA) for IL-13 responsive gene set derived from cytokine-treated air liquid interface (ALI) epithelial cultures. Genes were ranked by fold change between allergen challenge and baseline for HDM treated allergic asthmatics (AA). b. GSEA for IL-13 responsive gene set derived from cytokine-treated ALI epithelial cultures. Genes were ranked by fold change between allergen challenge and baseline for HDM treated allergic non-asthmatics (ANA). c. GSEA for IL-17 responsive gene set derived from cytokine-treated ALI epithelial cultures. Genes were ranked by fold change between allergen challenge and baseline for HDM treated AA. d. GSEA for IL-17 responsive gene set derived from cytokine-treated ALI epithelial cultures. Genes were ranked by fold change between allergen challenge and baseline for HDM treated ANA. e. Venn diagram describing the overlap between allergen challenge versus baseline differentially expressed genes between AA and ANA defined by raw p value less than 0.05. f. Venn diagram describing the overlap between allergen challenge versus baseline and allergen challenge versus diluent challenge differentially expressed genes between AA and ANA defined by raw p value less than 0.05. g. Heatmaps describing the DEGs from allergen challenge vs baseline and allergen challenge vs diluent challenge that were shared between AA and ANA. The heatmap on the left illustrates the DEGs that were differentially expressed in the same direction in both AA and ANA and the right illustrates the DEGs that were differentially expressed in opposite directions in AA and ANA. h. Calculation of the epithelial IL-13 responsive gene score at baseline compared to allergen challenge BAL PC1 from figure 2d for HDM challenged AA subjects i. Calculation of the epithelial IL-13 responsive gene score in diluent challenged samples compared to allergen challenge BAL PC1 from figure 2d for HDM challenged AA subjects j. Calculation of the epithelial IL-13 responsive gene score in allergen challenged samples compared to allergen challenge BAL PC1 from figure 2d for HDM challenged AA subjects k. Gene Ontology (GO) enrichment analysis for genes differentially expressed at baseline between type 2 high subjects (defined as PC1 > 0) and type 2 low subjects (Defined as PC1 < 0) for subjects challenged with HDM l. GO enrichment analysis for genes differentially expressed in allergen challenge samples between type 2 high subjects (defined as PC1 > 0) and type 2 low subjects (Defined as PC1 < 0) for subjects challenged with HDM

### Allergen challenge induces systemic clonally expanded airway Th2 cells

As expected, we have thus far shown allergen challenge induced stereotypical signs of local type 2 inflammation in the airways, particularly in individuals with indications of elevated type 2 inflammation at baseline. T helper 2 (Th2) cells induce and amplify type 2 inflammation. However, we were not able to clearly identify airway Th2 cells by CyTOF based on their selective expression of chemokine receptors (Strazza & Mor 2017; Lloyd & Hessel 2010). To further resolve the heterogeneity in the airway Th cell compartment, we performed single cell RNA sequencing (scRNAseq) and single cell TCR sequencing (scTCRseq) on a subset of AC and DIL samples from HDM-challenged individuals [Table 2]. To maximize cell yield and sequencing depth in T cells, we developed a sorting strategy to enrich CD4+ T cells from frozen BAL samples which were then pooled and encapsulated for single cell sequencing in a multiplexed fashion [Figure S2]. Using reference free genetic demultiplexing, this strategy clearly resolved CD4+ T cell subsets including Th1 cells expressing *CXCR3*, *TNF*, and *IFNG*; undifferentiated and naive cells expressing *CCR7*, *SELL*, and *IL7R*; T regulatory cells (Treg) expressing *FOXP3*, *TNFRSF18*, and *IL2RA*; cytotoxic-like cells expressing *GZMB*; interferon exposed cells expressing multiple interferon response genes such as *IFI6*; cycling cells marked by *MKI67*; and Th2 cells expressing *GATA3*, *IL5*, *IL13*, and *IL4* [Figure 4 a-c] (Hartoularos et al. 2023).

Allergen challenge in some subjects skewed the T cell landscape from one predominantly composed of Th1 cells to one predominantly composed of undifferentiated/naive cells [Figure 4d-e]. In addition, cells in the Th2 cluster were nearly exclusively found in AC samples. Since the prevalence of undifferentiated/naive cells in AC samples suggested an allergen-induced influx of cells from the systemic circulation, we calculated the single-cell gene set enrichment analysis score for the gene set defining human T resident memory (Trm) cells [Figure 4f] (Kumar et al. 2017). Cells in the Th1 cluster had the highest average Trm score, while the undifferentiated/naive cluster had the lowest Trm score as expected. Interestingly, the Th2 cluster displayed an intermediate phenotype, with Trm scores significantly lower than the Th1 cluster. This score stratification likely indicates that the undifferentiated/naïve cells and Th2 cells are more recent entrants into the lung microenvironment, whereas many of the Th1 cells are resident in the lung and present regardless of recent allergen challenge.

We also performed scTCRseq to evaluate the clonal response to allergen challenge within the CD4+ T cell compartment. We defined an expanded T cell clone as a group of two or more cells expressing the same T-cell Receptor Alpha (*TRA)* and T-cell Receptor Beta (*TRB*) sequences. As expected, expanded clones were private to each individual. Across individuals, they were present in both AC and DIL samples [Figure 5a], and at both day 1 and day 7 time points [Figure 5b]. The Th1, Th2, and Treg clusters all contained expanded clones. Both the fraction of T cells in each cluster belonging to expanded clones [Figure 5b] and the average number of cells within each expanded clone [Figure 5c] were higher in day 7 samples compared to day 1 samples. These expanded clonal characteristics in the differentiated clusters combined with a lack of clonal expansion in the undifferentiated/naive cluster strongly indicated allergen-driven clonal expansion in the Th1, Th2, and Treg clusters.

### Airway-infiltrating Th2 cell clones are allergen-responsive and present systemically

To test whether the expanded T cell clones were allergen responsive, we employed an activation induced marker (AIM) assay (Bacher et al. 2016; Bacher & Scheffold 2013). BAL samples were incubated *ex vivo* with HDM extract or PBS vehicle for 8 hours, then stained with the TotalSeq-C antibody cocktail for cellular indexing of transcriptomes and epitopes (CITE-Seq) analysis including both scRNAseq and scTCRseq of CD4+ T cells enriched as before by flow cytometry. The clusters identified by scRNAseq after these treatments had a distinct organization as compared to the samples directly assayed without stimulation [Figure 5d]. After stimulation, the data were primarily organized around activation status rather than cytokine expression profile [Figure 5e]. Within the clusters with high CD69 expression, the cluster which we call “*NR4A1 High 1*” contained the highest expression of *CD40L* mRNA and protein, and the TCR signal response gene *NR4A1*, indicating that this cluster contains cells receiving TCR-specific activation signals [Figure 5e]. FoxP3 expressing cells also formed multiple distinct clusters. The “Activated Treg” cluster contained cells with the highest expression of *IL2RA* mRNA and the canonical Treg activation marker protein CD137, indicating the presence of allergen-responsive Tregs [Figure 5e]. Importantly, T conventional and Treg cells within these clusters increased in prevalence in HDM stimulated samples compared with vehicle control samples [Figure 5e], as did the intensity of their activation marker expression [Figure 5f], further indicating their allergen-specific activation.

We then leveraged our scTCRseq analyses to trace T cell clones found in the AIM assay back to the directly assayed dataset, where cells within each clone were generally restricted to a single cluster with characteristics of recognizable differentiated T cell phenotypes [Figure S3]. We identified the dominant direct assay phenotype for each T cell clone, then annotated the AIM dataset by grouping the corresponding clones by these direct assay phenotypes [Figure 5g]. Th2 clones were distributed across multiple AIM assay clusters, with a minority of cells contained within activated clusters, mainly the *NR4A1* High 1 cluster [Figure 5g]. *NR4A1* High 1 cluster Th2 clones displayed the highest expression of *NR4A1* and all three major Th2 cytokines *IL4*, *IL5*, and *IL13* [Figure 5h]. Th1 clones occupied all of the non-Treg clusters in the AIM analysis, with the strongest representation among activated cell clusters [Figure 5g]. Th1 clones in the activated clusters had the highest expression of *IFNG*, *TNF*, *IL2*, and *NR4A1* [Figure 5i]. Treg clones retained FOXP3 expression in the AIM assay and were enriched in the Activated Treg cluster defined by upregulation of CD137 (Saggau et al. 2021), as well as increased *TGFB1*, *NR4A1*, *IL10*, and *EBI3* expression [Figure 5g&j]. These data indicate the stability of the Treg phenotype and show that many of the Treg clones detected in the BAL were indeed allergen responsive. Undifferentiated/naive clones were primarily found in the unactivated clusters in the AIM dataset, and those few cells found in activated clusters did not have strong cytokine or NR4A1 expression, supporting their unreactive, undifferentiated state [Figure 5k].

These data demonstrate that clones of Th1, Th2 and Treg cells from the airways responded to HDM with gene expression patterns consistent with allergen-specific TCR signaling. Yet only Th2 cells were specifically overrepresented in AC compared with DIL samples (Figure 4d&e), suggesting that allergen-responsive Th2 cells may be recruited to the lung upon allergen exposure. Therefore, we sought to determine whether these clones exhibited indications of systemic activation and expansion. Bulk TCR sequencing from RNA isolated from peripheral blood revealed that across all samples many of the TCRs found in both the directly assayed samples and the AIM assay were present in the blood both before (visit V2) and after (V3) *in vivo* allergen challenge [Figure 5 l-n & Table 3]. In two day 7 samples with high type 2 inflammation, indicated by PC1 (ACE-034 and ACE-036), *TRA* sequences from airway Th2 clones were enriched in the blood during visit V3 compared to V2 [Figure 5l]. Interestingly, the most enriched *TRA* sequences in the blood matched with Th2 clones with an activated phenotype in the *ex vivo* allergen stimulation assay [Figure 5l]. We did not observe this trend consistently in *TRA* sequences from Th2 clones that were not activated in the *ex vivo* stimulation assay, nor those from Th1 or Treg clones [Figure 5l-n]. We observed a very similar pattern in the blood frequencies of the corresponding *TRB*s from these same airway T cell clones [Figure S4]. These results demonstrate that allergen-responsive Th2 cells are present but rare in the airways even in the setting of allergic asthma, but they are easier to detect at day 7 after allergen exposure in individuals with high levels of type 2 inflammation. Further, antigen-responsive Th2 cells are present in the blood prior to allergen exposure, airway allergen challenge induces systemic expansion that is not limited to the local lung environment.

**Table 2.** 10X scRNAseq Samples Assayed.

| ID | V3 Time Point | Diagnosis | Allergen | AC/DIL |
| --- | --- | --- | --- | --- |
| ACE-28 | 1 day | AA | HDM | AC only |
| ACE-34 | 7 day | AA | HDM | both |
| ACE-36 | 7 day | AA | HDM | both |
| ACE-37 | 1 day | AA | HDM | both |
| ACE-42 | 1 day | NANA | Cat dander | both |
| ACE-43 | 1 day | AA | HDM | both |
| ACE-44 | 1 day | AA | HDM | both |
| ACE-49 | 1 day | AA | HDM | both |
| ACE-71 | 1 day | AA | HDM | both |

**Figure 4.**
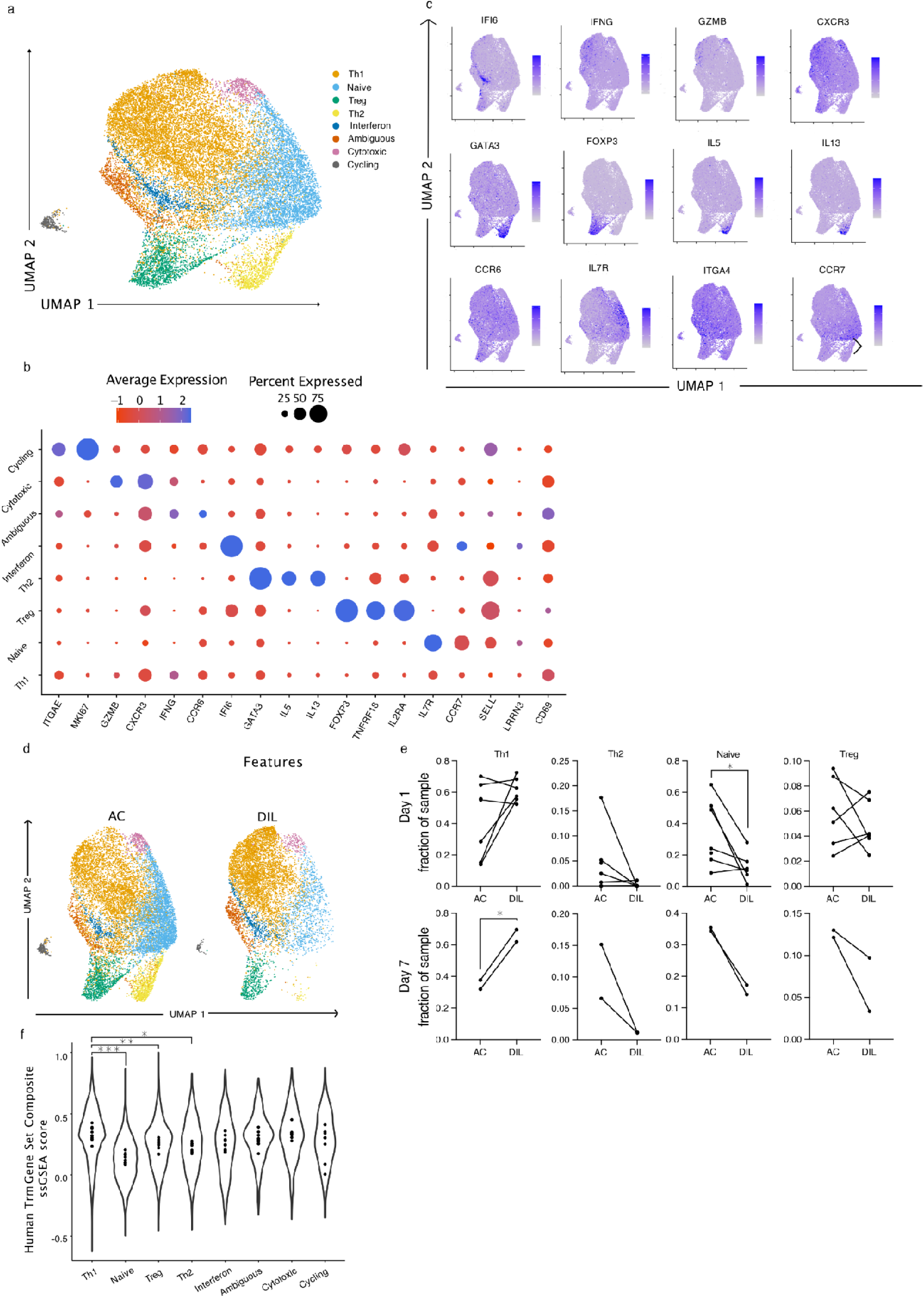
scRNAseq on enriched CD4+ T cells resolves allergen induced Th2 Cells a. UMAP illustrating CD4+ T cell subsets identified by unbiased clustering of scRNAseq from frozen samples enriched for CD4+ T cells via flow cytometry b. Dot plot illustrating marker gene expression in the clusters identified in (a) where color indicates intensity of expression and dot size represents percentage of cells within that cluster expressing a given gene c. Feature plots for marker gene expression in the UMAP space defined in (a) and darker blue color indicates increased intensity of given gene expression d. UMAP split by V3 sample treatment of either allergen challenge or diluent challenge with each cell color coded by unbiased clustering assignment e. Allergen challenge versus diluent challenge representation of each cluster identified in (a) as a fraction of sample identified by the given cluster. Lines indicate paired samples obtained from the same subject. Statistics displayed determined by paired t-test between AC and DIL samples (*, p<0.05) f. Single cell gene enrichment scores for a tissue residency gene program defined by (Kumar et al. 2017) plotted for each cluster identified in (a) for all samples assayed. Each dot represents the mean value of the gene score for all cells in that cluster belonging to a given subject. The violin plot illustrates the population distribution for all cells in the identified cluster. Statistics displayed are determined by paired one-way ANOVA with post-hoc testing of differences between every pair of clusters with p-value adjustment for multiple hypothesis testing. For clarity, a subset of calculated p-values are displayed. (*,p<0.05;**,p<0.01;***,p<0.001).

**Figure 5.**
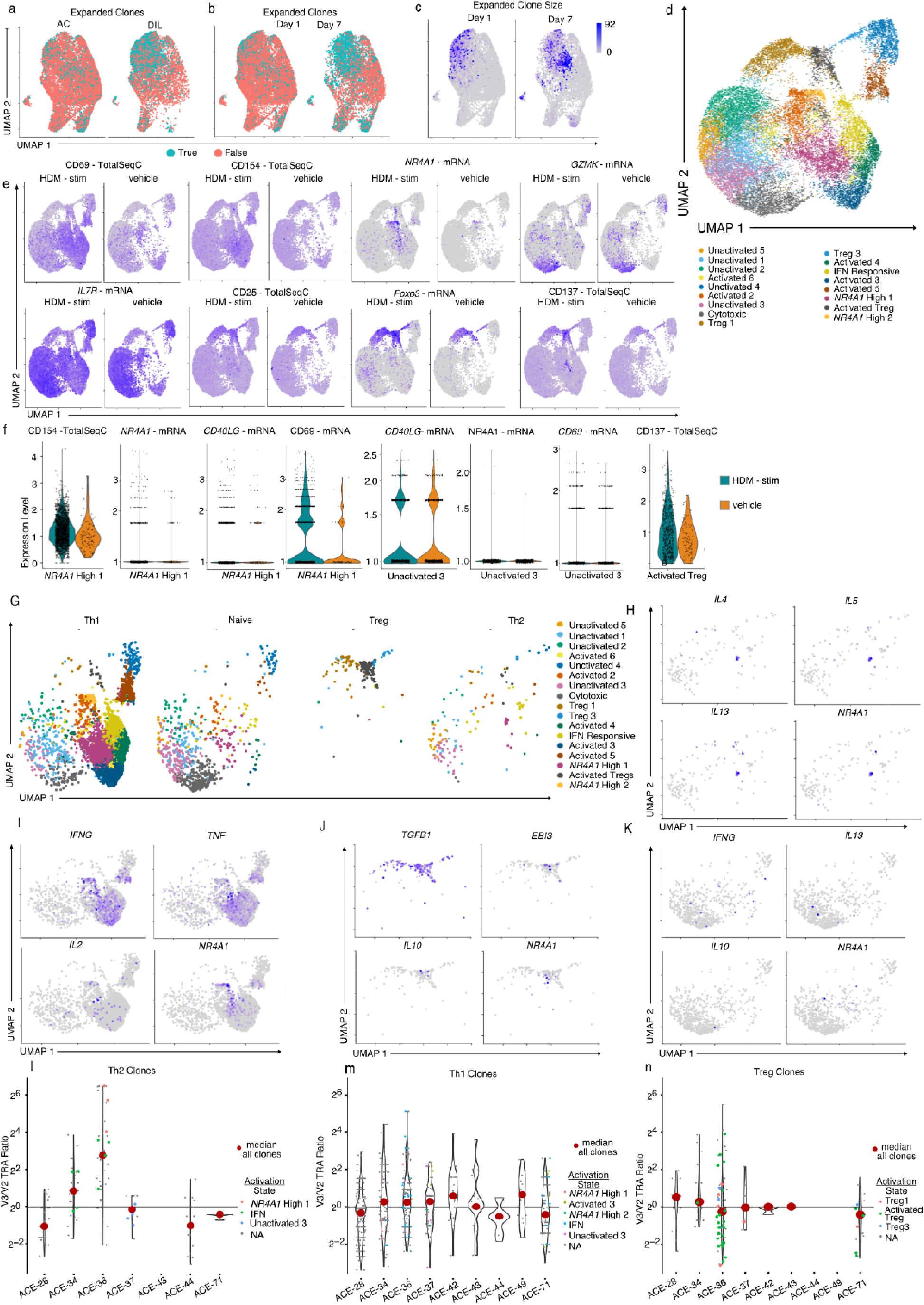
Expanded Th2 clones are allergen reactive and expand in the blood post-challenge a. UMAP of CD4+ T cells identified in the directly assayed data set described in Figure 4. Individual cells are color coded by their status as part of an expanded T cell clone, defined by 2 or more cells containing the exact same *TRA* and *TRB* pair. The plot contains all samples assayed and is split by the treatment condition of the sample. b. UMAP of CD4+ T cells identified in the freshly assayed data set described in Figure 4. Individual cells are color coded by their status as part of an expanded T cell clone, defined by 2 or more cells containing the exact same *TRA* and *TRB* pair. The plot contains all samples assayed and is split by the time point of the sample. c. UMAP of CD4+ T cells identified in the freshly assayed data set described in Figure 4. Individual cells are color coded by how many other cells belong to their T cell clonotype, defined by 2 or more cells containing the exact same *TRA* and *TRB* pair. The plot contains all samples assayed and is split by the time point of the sample. (d-k)Activated induced mark assay where BAL aliquots were thawed and stimulated for 8 hours with HDM extract and then sorted by flow cytometry prior to analyzing gene expression and TCR sequences by scRNAseq d. UMAP of CD4+ T cells from the AIM assay. Individual cells are color coded by their unbiased cluster designation. e. Feature plots of key marker gene/protein expression in the AIM assay split between the *ex vivo* treatment conditions. Gene expression detected via direct sequencing of the mRNA molecule is denoted with “-mRNA”. Protein expression detected via oligo tagged antibody binding to the marker of interest is denoted with “-TotalSeqC”. Darker blue indicates increased expression of that gene/protein. f. Gene/protein expression of key TCR activation induced genes in cluster designated “activated 1” and cluster designated “unactivated 3”. Each dot indicates the expression level of the given gene/protein for a single cell in that cluster and *ex vivo* treatment. All samples are included and split by *ex vivo* treatment. Violin plot captures the distribution for a given gene/protein in that cluster and *ex vivo* treatment group. g. UMAP as defined in d, but split by the phenotype of each clonotype in the directly assayed data set. TCR sequences were used to barcode the cells present in both the directly assayed aliquots and the *ex vivo* stimulated AIM assay. Clonotypes were assigned a phenotype from the cluster most represented in that clonotype from the directly assayed data set. h. Feature plots illustrating type 2 cytokine and activation induced gene expression in AIM clones containing TCR clonotypes defined as Th2 in the directly assayed data set i. Feature plots illustrating type 1 cytokine and activation induced gene expression in AIM clones containing TCR clonotypes defined as Th1 in the directly assayed data set j. Feature plots illustrating regulatory T cell cytokines and activation induced gene expression in AIM clones containing TCR clonotypes defined as Tregs in the directly assayed data set k. Feature plots illustrating type 1, type 2, and regulatory T cell cytokines and activation induced gene expression in AIM clones containing TCR clonotypes defined as undifferentiated/naive in the directly assayed data set (l-n) TCR sequencing was performed on bulk RNA isolated from peripheral blood at clinical visits 2 and 3. *TRA* prevalence was determined by the percent of total TCR reads a given alpha chain occupied. V3/V2 enrichment was determined by the ratio of prevalence for a given *TRA* at visit 3 divided by the prevalence for the same *TRA* at visit 2. l. Enrichment of *TRA* sequences associated with clonotypes identified as Th2 clones in the directly assayed data set. Select clones that were also identified in the AIM assay are color coded by the most prominent cluster from that assay. m. Enrichment of *TRA* sequences associated with clonotypes identified as Th1 clones in the directly assayed data set. Select clones that were also identified in the AIM assay are color coded by the most prominent cluster from that assay. n. Enrichment of *TRA* sequences associated with clonotypes identified as Treg clones in the directly assayed data set. Select clones that were also identified in the AIM assay are color coded by the most prominent cluster from that assay.

**Table 3.** TCRs in directly assayed scRNAseq data set are broadly found in the blood before and after challenge.

| Challenge Condition | Total BAL TRAs | Total BAL TRBs | Expanded BAL TRAs | Expanded BAL TRBs | Expanded BAL TRAs in Blood V2 | Expanded BAL TRBs in Blood V2 | Expanded BAL TRAs in Blood V3 | Expanded BAL TRBs in Blood V3 |
| --- | --- | --- | --- | --- | --- | --- | --- | --- |
| AC | 7964 | 8183 | 1069 | 1058 | 413 | 623 | 431 | 640 |
| DIL | 3831 | 3819 | 956 | 919 | 336 | 518 | 283 | 453 |

### Allergen induces monocyte derived populations expressing T cell chemoattractants

When we enriched CD4+ T cells for these analyses, we also collected other cells in the airway samples for parallel analysis by scRNAseq. However, the abundant granulocytes, particularly eosinophils and neutrophils, presented a technical challenge for the generation of high quality scRNAseq data (Lei et al. 2021; Martin et al. 2019). To address this issue, we employed a flow cytometry strategy to simultaneously sort CD4+ T cells in one tube, and a pool of all other leukocyte populations that were not CD4+ T cells, neutrophils, or eosinophils in another tube [Figure Sx z]. In the latter pool, we identified 9 cell type clusters by unbiased nearest neighbor clustering, including both myeloid and lymphoid immune cells [Figure 6 a-c]. Among the 4 myeloid clusters, 3 represented a spectrum of macrophages and 1 represented dendritic cells [Figure 6a-c]. Many of the dendritic cells express markers associated with an infiltrating monocyte origin (*CD1A*, *FCER1A*, and *CXCR4*) [Figure 6 a-c] (Collin & Bigley 2018). The macrophage clusters all expressed *MARCO* and *MRC1*, but unbiased analysis indicated that *CXCR4, FABP4, CD14, VCAN, CCL2,* and *CCL13* were expressed in patterns unique to each cluster [Figure 6b-c]. Of particular interest, the *VCAN and CCL2 high macrophages* demonstrated hallmarks of M2-like or alternatively activated macrophages that are associated with allergic disease [Figure 6b-c] (Saradna et al. 2018). These VCAN and chemokine high macrophages also appear to be monocyte-derived due to the high expression of *CD14* [Figure 6 b&c]. Of the other two major macrophage clusters, both were defined by *FABP4, MARCO*, and *MRC1,* suggesting these may represent tissue resident alveolar macrophages with one cluster expressing *FABP4* at a higher level and the other low for *MRC1* expression [Figure 6b&c] (Liang et al. 2019). The final macrophage clusters identified are interferon responsive with high expression of *SIGLEC1*, a gene shown to be induced by viral infection of macrophages [Figure 6 b&c](Herzog et al. 2022). The final myeloid cluster present closely associated with other macrophage clusters, contained very few cells, and was marked by *CYP2S1* and *ITGA3* expression [figure 6 b&c]. In the first analysis of all cells, we delineated a clear B cell cluster marked by *MS4A1* and *JCHAIN* expression, and a mixed cluster of cytotoxic lymphocytes containing *CD8A, TRDC*, and *NKG7* expressing cells [Figure 6 b&c]. We further resolved this cytotoxic cell cluster by sub-clustering and marker gene analysis [figure 6d-f]. Among the resultant sub-clusters, 3 contained CD8+ T cells (one mixed with macrophages) primarily delineated by the expression level of IL32 [Figure 6 d-f]. Two clusters contained NK cells and gamma delta T cells that could not be fully resolved, with unbiased analysis dividing these into two clusters based in part upon *GNLY* expression[figure 6 d-f].

We then sought to determine how allergen challenge changed this landscape. In contrast to the T cell compartment, day 1 samples did not yield any clear statistically significant changes and the only statistically significant change in the day 7 samples was an increase in the combined mixed lymphocyte population [Figure 7 a&b]. As cell population changes were not evident, we next evaluated if gene expression programs were influenced by allergen challenge. Strikingly, unbiased evaluation of DEGs indicated that chemokine gene programs were strongly induced in several of the major clusters [Figure 7 c&d]. Notably, in the monocyte derived M2-like macrophages and the monocyte derived dendritic cells *CCL3* (Log2FC 71.5 & 146.8 respectively), *CCL17* (Log2FC INF & INF respectively), *CCL22* (Log2FC 121,7 & 110 respectively), and *CCR1* (Log2FC 14.8 & 35.8 respectively) were induced by allergen challenge [Figure 7 c&d]. Further, *CCR5* and *CCR6* were induced on the monocyte derived dendritic cell population [Figure 7 c& d]. In a similar pro-inflammatory fashion, *CCL5* was induced by allergen challenge in the mixed lymphocyte population [Figure 7 c&d]. In opposition to these allergen induced changes, in the tissue resident alveolar macrophage population CCL18 was reduced by allergen challenge [Figure 7 c&d]. As these chemokines have been shown to have important chemotactic properties for Th2 cells, we then sought to see if similar programs were induced by allergen challenge in the CD4+ T cell compartment [Figure 7 e](Pilette et al. 2004). In contrast to the induction of chemokine receptors in macrophage populations we did not observe any change in *CCR4* or *CCR5* induced by allergen [Figure 7 e]. Further, the distribution of these receptors was generally broad [Figure 7 e]. In line with previous work, expression of *PTGDR2* (CRTH2) and lack of CCR6 best identified Th2 cells’ chemokine receptor profile [Figure 7e & Figure 4 b&c] (Hirai et al. 2001). Interestingly, Tregs exclusively expressed *CCR8* and the expression of this receptor was higher in the diluent challenged samples, connecting Tregs strongly to CCL18 expression by the tissue resident alveolar macrophages [Figure 7 c-e]. *CCL17* (TARC) and *CCL22* have been shown to be important mediators of type 2 inflammation in asthma by recruiting *CCR4*+ Th2 cells (Pilette et al. 2004). We compared the presence of the mo-DC and monocyte derived macrophage populations expressing *CCL17* and *CCL22* with the Th1 and Th2 populations from the allergen challenged samples [Figure 7 f]. Surprisingly, both of these populations correlated negatively with Th2 cells, and both of them correlated positively with Th1 cells [Figure 7 f]. This result suggests that macrophage and monocyte derived populations in the lung are important to define the chemokine signature shortly after allergen exposure. However, chemokines like *CCL5*, *CCL17*, and *CCL22* may not have Th2 specific effects in the lung due to wide expression of the receptors *CCR5* and *CCR4* on various T cell subsets.

**Figure 6.**
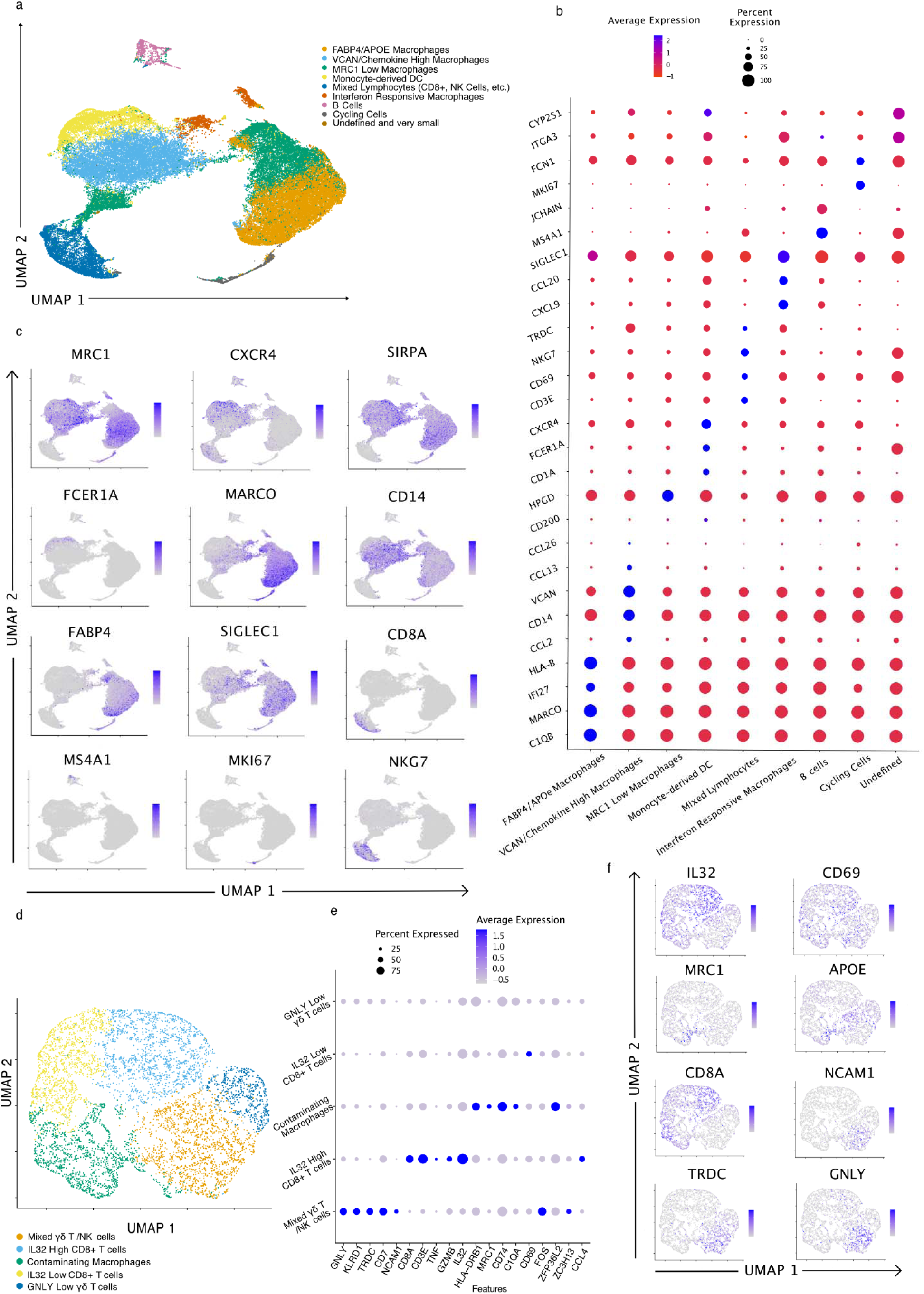
scRNAseq defines non-granulocyte, non-CD4+ T cell populations Frozen BAL aliquots were thawed and sorted by flow cytometry to remove neutrophils and eosinophils. CD4+ T cells were separately assayed as shown in figure 4 & 5, the rest of the cells in the sample were assayed scRNAseq. a. UMAP showing dimensionally reduced visualization of scRNAseq data. Each dot represents a cell color coded by the cluster identified through unbiased nearest neighbors clustering. All samples displayed. b. Dot plot indicating key marker gene expression. Dot size indicates percent of cells in a given cluster expressing a given gene and color indicates the intensity of expression. c. Feature plots visualizing key marker gene expression displayed in UMAP space defined in (a). Increasingly dark blue color indicates increased expression of the indicated gene (d-f) The mixed cytotoxic lymphocyte cluster from the whole data set was subset and dimensional reduction and clustering analyses were performed to further resolve the populations. d. UMAP showing dimensional reduced visualization of sub-clustered cytotoxic lymphocytes. Each dot represents a cell color coded by the new higher resolution unbiased nearest neighbors clustering. All samples are displayed. e. Dot plot indicating key marker gene expression. Dot size indicates percent of cells in a given cluster expressing a given gene and color indicates the intensity of expression. f. Feature plots visualizing key marker gene expression displayed in UMAP space defined in (d). Increasingly dark blue color indicates increased expression of the indicated gene

**Figure 7.**
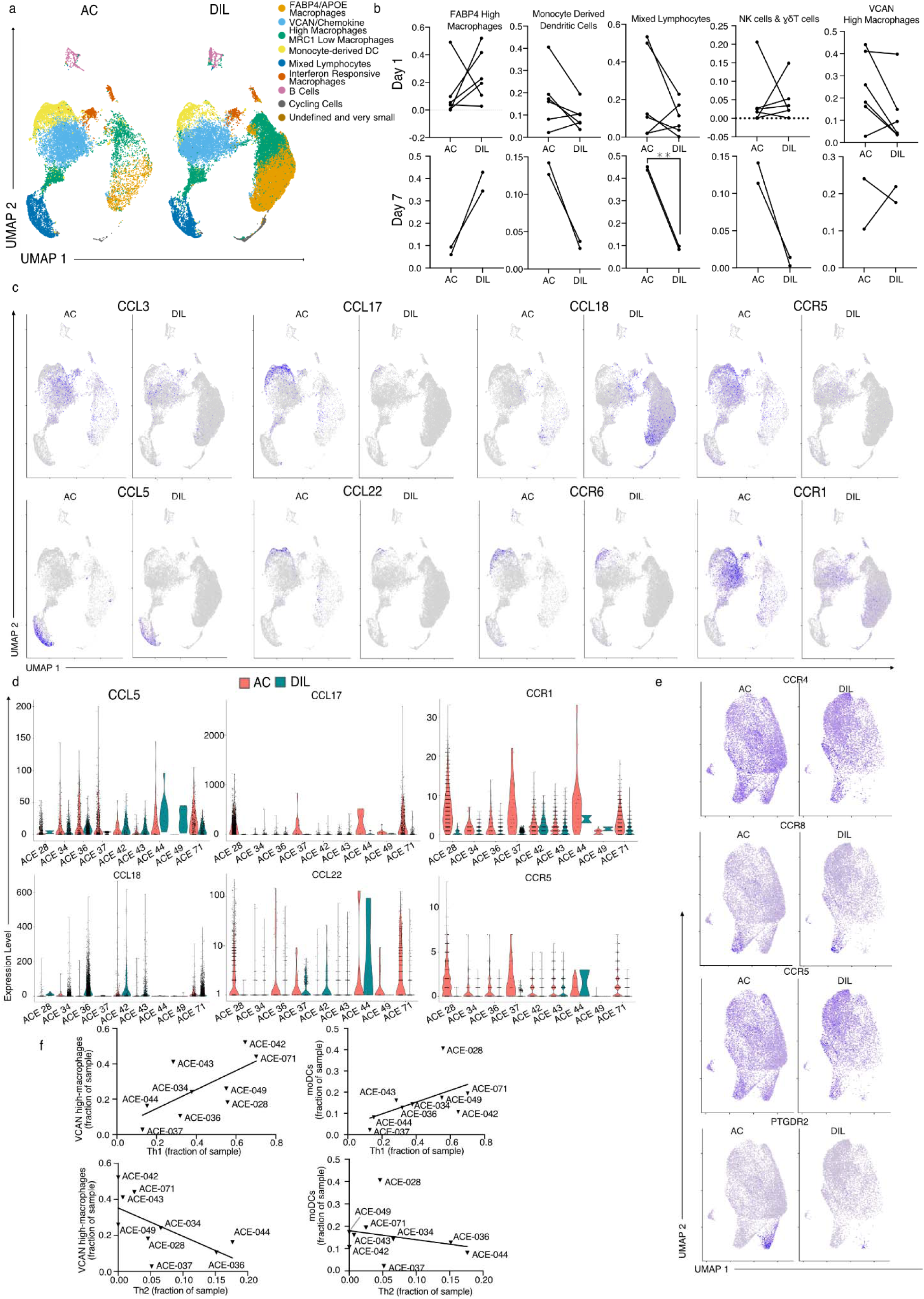
Allergen induces T cell recruiting chemokine expression in monocyte derived populations Single cell analysis defined in figure 6 is analyzed for differences in cellular composition and gene expression and is connected to gene expression and cellular composition in the CD4+ T cell data set defined in figure 4. a. UMAP dimensionally reduced plot from figure 6a split by V3 sample treatment. b. Allergen challenge versus diluent challenge representation of each cluster identified in (a) as a fraction of sample identified by the given cluster. Lines indicate paired samples obtained from the same subject. c. Feature plots split by allergen challenge and diluent challenge of key chemokine and chemokine receptor genes identified as differentially expressed by Wilcoxon Rank Sum test. d. Gene expression of differentially expressed key chemokine and chemokine receptor genes broken down by subject and sample treatment. *CCL5* is shown for the mixed cytotoxic lymphocyte cluster. *CCL17* and *CCL22* are shown for the mo-DC cluster. *CCR1* and *CCR5* are shown for the monocyte derived macrophage cluster. *CCL18* is shown for the tissue-resident alveolar macrophage cluster. Each dot represents a single cell and the violin plot illustrates the population distribution. e. Feature plots indicating the expression of chemotactic receptors on the directly analyzed data set from figure 4, split by sample treatment and color intensity by gene expression. f. Correlation between monocyte derived populations and Th1 and Th2 subsets defined in the directly analyzed data set from figure 4. The plotted relationship line represents linear regression of the two variables. Statistics displayed determined by paired t-test between AC and DIL samples (**, p<0.01)

## Discussion

In this study, we combined high dimensional cell analysis techniques to generate an integrated atlas of the human asthmatic response to allergen, incorporating epithelial cell gene expression, broad cellular landscape changes, and specific changes within inflammatory cells. The scRNA-seq and CyTOF performed here primarily with BAL confirms that increased cellular inflammation, IL-13 responsive gene expression in myeloid and epithelial cells, and Th2 cells are key nodes in asthmatic airway responses to allergens. These findings complement prior analyses of BAL from unchallenged asthmatic subjects (Seumois et al. 2020), and of brushings of the lower airway mucosa with and without allergen challenge (Alladina et al. 2023; Rahimi et al. 2026). Integrated analysis implicates allergen-induced chemoattractant circuits in the recruitment of Th2 cells and other inflammatory cells to the airways. Importantly, we identify allergen-reactive Th2 cell clones that already exist systemically, then expand and infiltrate the airways upon local allergen exposure. Taken together, our data demonstrate that allergen exposure induces coordinated changes to the airway microenvironment through local changes in inflammatory gene expression and ingress of allergen-reactive Th2 cells.

Understanding the heterogeneity of the local lung response to allergen is critical, as the most detrimental aspects of asthma, such as the formation of mucus plugs, are localized in nature (Tang et al. 2022). Critical to this is identifying the inflammatory circuits that drive the recruitment of particular cell populations. Through unbiased differential gene expression analysis, we show that *CCL17* and *CCL22* are induced by allergen challenge within monocyte derived macrophage and dendritic cell populations. These chemokines induce cell recruitment through their shared receptor, *CCR4,* which is consistently expressed on Th2 cells (Pilette et al. 2004). However, our data indicate that these chemokines likely act broadly on the T cell compartment in allergen challenged human airways, as *CCR4* expression was promiscuous among BAL T cells. Th2 cell recruitment may be more specifically driven by the prostaglandin D2 receptor CRTH2, which was highly selectively expressed by Th2 cells. Our data also highlight other interesting aspects of homing receptor expression. *CCR5* was also broadly expressed on airway T cells. The Th17-associated chemokine receptor *CCR6* was sparsely expressed with little specificity for particular CD4+ T cell subsets outside of consistently low expression in Th2 cells. This is consistent with the poor *IL17A* and *IL17F* expression observed in this data set, and with the relatively low involvement of these pathways in allergic asthma (Melgert et al. 2007). In the Treg compartment, *CCR8* stands out as a specifically expressed chemokine receptor of likely functional relevance, as its ligand CCL18 was also produced by tissue resident alveolar macrophages. This circuit also plays an immunoregulatory role in mouse models of asthma (Jheng et al. 2023). Taken together, these data confirm prior studies in mouse and less detailed human studies, enhancing their relevance to human airway allergen responses and providing novel insights connecting local and systemic T cell responses.

This study, in the context of other recent work, clearly demonstrates local Th2 cell ingress to the airway upon allergen challenge in human allergic subjects. Allergen-reactive T cells have been detected in the peripheral blood ((Bacher et al. 2016) and BAL (Seumois et al. 2020) of allergic subjects. scRNAseq clearly discerns Th2 cells in allergic airways by their expression of the canonical transcription factor GATA3 and cytokines IL-4, IL-5, and IL-13 (Alladina et al. 2023; Rahimi et al. 2026; Seumois et al. 2020; and this study). We show that these Th2 cell populations contain subject-specific expanded clones that are more evident 7 days post allergen challenge compared to 1 day post challenge. A subset of these clones was allergen-reactive, present in the blood before airway allergen challenge, and also expanded in the blood repertoire after allergen challenge. These findings connect systemic allergen-reactive cells to elevated, allergen-driven type 2 inflammation in the lower airways, providing novel evidence in humans for a paradigm in which Th2 cells are produced systemically and recruited to the airways by chemoattractant and other inflammatory cues associated with atopic disease (Lambrecht et al. 2019).

The ability to link clonal Th2 cells which are present in blood and expanded in the lung after allergen challenge in asthma has important clinical implications. Circulating pathogenic cells may be a druggable target through inhibition of their recruitment to the lung. Indeed, whether dupilumab or other asthma biologics work by affecting tracking or survival of these clones is unknown. Furthermore, peripheral blood T cell clones could become a meaningful biomarker, and personalized immune targeting becomes feasible if specific pathogenic clonotypes can be identified and targeted, perhaps using peptide immunotherapy or TCR-informed allergen immunotherapy refinement. The value of such an approach would be its potential to induce immunological remission, which contrasts with current asthma therapies.

The kinetics of Th2 cell recruitment and expansion following airway allergen challenge have interesting implications for the functional impact of these cells at the airways. Eosinophils were early entrants upon allergen challenge, and monocyte derived populations and T cells expressing type 1 cytokines were also significant infiltrating populations in the airways. Th2 cells were particularly enriched at later time points following allergen exposure. Extended presence of Th2 allergen reactive cells in the airways may induce longer term tissue remodeling, eosinophil survival and chemotaxis by providing IL-5 directly at and within the airways. As long term airway remodeling is a critical part of asthma, our data highlight chemokine pathways that are essential to acute airway inflammation and the cellular basis for type 2 cytokines acting over longer time frames.

We integrated epithelial gene expression data with single cell level molecular and cellular analysis of allergen-induced airway inflammation to define a type 1 / type 2 axis across subjects. In type 2 high individuals, we observed clonally-related Th2 cells, at least some of which are allergen-reactive and present systemically. Future studies should address the spatial and temporal dynamics of allergen-specific Th2 cell responses in the lung and peripheral immune organs. Understanding allergen uptake and the trafficking of lung resident and/or infiltrating antigen presenting cells may reveal how allergen-specific Th2 cell prevalence is sustained over time and inform the prioritization and development of therapeutic strategies to minimize both acute and sustained inflammation in allergic asthma.

## Materials and Methods

### Human Subjects and Clinical Study

Human samples were collected as part of the Allergen Challenge for Evoked Phenotypes in Asthma (ACE Study, AADCRD-UCSF-01), which enrolled participants with asthma and healthy controls (NCT02230189). Participants were recruited through community-based advertising. The study aimed to provide a defined allergic stimulus to induce cellular and molecular events, facilitating the analysis of time-course changes. The primary focus was on subjects with stable or well-controlled asthma who were allergic to either House Dust Mite (HDM) or cat dander. A smaller number of non-asthmatic and non-allergic controls were also included. Inclusion criteria for participants with asthma included age between 18 and 50, self-reported prior physician-diagnosed asthma, pre-bronchodilator baseline Forced Expiratory Volume in 1 second (FEV1) greater than 75% of predicted, positive skin test reactivity (wheal ≥3mm) to house dust mite or cat allergen, and methacholine PC20 (provocative concentration causing a 20% drop in FEV1) less than 16 mg/mL. Healthy controls had no history of asthma or other lung diseases, negative skin test reactivity, FEV1 greater than 90% of predicted, and methacholine PC20 greater than 16 mg/mL. Exclusion criteria included current smoking, systemic steroid use in the six months prior to enrollment, inhaled steroid use within the previous month, and current indoor cat and/or indoor second-hand cigarette smoke exposure within the one month prior to the study. The study was approved by the University of California at San Francisco institutional review board, and written informed consent was obtained from all study participants before enrollment.

Samples were collected during three visits for participants. The initial visit involved characterization through history, spirometry, and quantitative allergen skin prick testing to determine safe doses for segmental allergen challenge. One week after the first visit, bronchoscopy was performed to obtain baseline bronchoalveolar lavage (BAL) from the left upper lobe (LUL) and airway brushings from the left lower lobe (LLL). Then diluent was administered in a segment of either the right middle lobe (RML) or right upper lobe (RUL, selected randomly) and allergen was administered in either the RML or RUL (the lobe which did not receive diluent). During the third visit, bronchoscopy was performed to collect BAL and airway brushings from the challenged segments. This study is largely similar to those previously conducted (Cho et al. 2016). It is important to note that the sample size represents a subset of the entire study, and the selection criteria for these samples were defined separately. The study received support from the NIH/NIAID grant 5U19AI077439 and was registered on clinicaltrials.gov under the identifier NCT02230189.

### Skin Prick Testing

Qualitative skin prick allergen reactivity testing was performed using the Multi-Test II skin test applicators. The applicator was carefully removed from the Dipwell Tray and pressed into the forearm skin of the subject with sufficient pressure to allow adequate penetration of the points. The subject was monitored during reaction incubation, and the histamine and saline negative controls were read at 15 minutes. The allergen results were read at 20 minutes by outlining each wheal with a marking pen. A wheal diameter of 3mm in diameter was considered a positive reaction. Based on the results of the qualitative skin prick test, a quantitative skin prick test was done using the Morrow Brown Disposable Skin Testing Needles. Serial dilutions of either cat dander or HDM were chosen based on the qualitative skin test. Using the Disposable Skin Testing needle a drop of each dilution was applied to the skin and then a prick with the needle was administered in the middle of the droplet. After 20 minutes, the reaction wheal were read in the same way as before. The lowest concentration to elicit a wheal reaction great than 3 mm in diameter was considered the “Threshold concentration”.

### Segmental Allergen Challenge

To perform segmental allergen challenge (SAC) at the V2 bronchoscopy, 2 mL of either diluent or allergen were administered into distinct lobes of the lung using a protocol similar to that used in a previously published study(Cho et al. 2016) . Diluent was administered in a segment of either the right middle lobe (RML) or right upper lobe (RUL, selected randomly) and allergen was administered in either the RML or RUL (the lobe which did not receive diluent). For safety, a test dose of allergen was administered first. For HDM, the test dose consisted of 1/30th the threshold allergen concentration determined from skin prick testing. If after at least two minutes, there was no evidence of mucosal inflammation a second larger dose of allergen was administered. This dose consisted of 2 mL of HDM allergen at 1/3rd the threshold allergen concentration (maximum cat dose = 740 BAU). For cat allergen, the test dose also consisted of 1/30th the threshold allergen concentration determined from skin prick testing. If after at least two minutes, there was no evidence of mucosal inflammation, the full dose consisted of 2 mL of cat allergen at the full threshold allergen concentration (maximum cat dose = 1,000 BAU). V3 BAL samples were collected from the right upper lobe and the right middle lobe via 3, 50 mL installations in each lobe.

### Bronchoalveolar lavage (BAL)

Bronchoalveolar lavage (BAL) was performed during bronchoscopy by instilling three 50-mL aliquots of sterile saline (37 °C) into a segmental bronchus (total 150 mL). Fluid was aspirated using wall suction after each aliquot until return slowed or ceased and collected into lidocaine-free specimen traps kept on ice. Typical returns were 60–90 mL. BAL fluid was filtered through two-ply gauze into 50-mL conical tubes.

### Cytospins

The cells were counted in the BAL fluid and 10 mL of BAL fluid was normalized to a cellular concentration such that it will yield 40,000 to 50,000 cells per cytocentrifuge slide. 60 µL of cell suspension was transferred into four shandon cytopsin funnels and centrifuged at 500 rpm for 5 minutes. Slides were stained using the Shandon Diff-quick kit. Slides were subsequently counted and evaluated by light microscopy.

### mRNA Sequencing from Epithelial Brushings

Bronchial epithelial brushings were performed during bronchoscopy using single-use cytology brushes (3 mm × 11 mm; ConMed Corporation, Utica, NY; catalog no. 149) deployed at segmental and sub-segmental bronchi. Collected epithelial cell brushing remaining after cytospin preparation and cell counts were spun at 500 rpm for 5 minutes. The supernatant was discarded and the cells were disrupted by adding 600 µL of QIAzol. Samples were vortexed for 1 minute to ensure complete lysis of cells. Samples were stored at -80°C. Total RNA was extracted using the miRNeasy Mini Kit (Qiagen, Hilden, Germany) according to the manufacturer’s protocol. RNA concentration and purity were assessed by A260/A280 ratio using a Nanodrop spectrophotometer (Thermo Fisher Scientific). RNA integrity was evaluated using an Agilent Bioanalyzer (Agilent Technologies).

### Sample Storage and CyTOF

BAL cells were counted using Turks solution and hemocytometer and processed for cytometry by time-of-flight (CyTOF) staining. Red blood cells were lysed using red blood cell lysis buffer for 5–6 min, followed by centrifugation and washing twice with PBS (CyPBS). Cells were counted and 3 × 10 cells were aliquoted per sample and washed twice in staining buffer (CyFACS). Cells were incubated with a metal-conjugated antibody cocktail for 30 min at room temperature (50 µL staining solution per 3 × 10 cells). After staining, cells were washed twice with cell staining media (CSM) and viability was assessed. Cells were then resuspended at 2 × 10 cells/mL and incubated with cisplatin (2 µM; Cell-ID Cisplatin, DVS Sciences) for 5 min at room temperature for viability labeling, followed by quenching with excess CSM. Cells were fixed with 2% paraformaldehyde and stored at 4 °C overnight. Following fixation, cells were washed in CSM, permeabilized with saponin-containing buffer for 45 min on ice, and washed again prior to nucleic acid staining. DNA was labeled using an iridium intercalator (1:5000 dilution in intercalation buffer) for 20 min at room temperature. Cells were subsequently washed with CyFACS buffer and MilliQ water prior to acquisition on a CyTOF mass cytometer. Samples not used for CyTOF were resuspended in 10% DMSO, 10% FCS, RPMI and stored at -80°C.

### Bulk TCR Sequencing from Peripheral Blood

Peripheral Whole blood was collected on clinical visits 2 and 3 into PAXgene tubes to collect stabilized RNA. RNA was subsequently isolated using the Qiagen PAXgene kit and stored at - 80°C. RNA quality was assessed by the agilent bioanalyzer and 14 out of 18 had RNA integrity (RIN) scores > 8, 2 out of 18 had RIN scores > 7, and 2 out of 18 had RIN scores ≥ 6.5. Isolated RNA was used as input into the Takara SMARTer Human TCR ab Profiling kit for next generation sequencing library preparation. These libraries sequenced on the NovaSeq6000 SP using PE-150 reads. FASTQs generated were analyzed using the Cogent NGS Immune Profiler software provided by Takara to generate TRA and TRB sequences present and the fraction contained within the sample.

### scRNAseq and scTCRseq from BAL

Frozen BAL aliquots were thawed at 37°C and washed with RPMI-1640, 20% FCS, 25 µg/mL DNase I. Samples were then passed through a 40 µm filter and washed 1X with 2% FCS, 1 mM EDTA PBS. The samples were then stained with Invitrogen™ eBioscience™ Fixable Viability Dye eFluor™ 780, CD4 PE-Cy7, CD45 ef450, CD3E APC, CD20 FITC, CD56 FITC, CD14 FITC, CD19 FITC, CD16 FITC, CD15 AF700, CD24 PE, and CD206 BV605. Cells were then resuspended in 2% FCS, 25 mM HEPES (pH 7.0), 1 mM EDTA PBS for sorting on the Aria Fusion S854 with biosafety cabinet and aerosol control. Samples were sorted to remove granulocytes (eosinophils and neutrophils) and separate CD4+ T cells from the remaining populations. CD4+ T cell samples from 4 distinct individuals at a time were mixed and resuspended at 1000 cells/µL. These mixed samples were then run on a single lane of a 10X chip for GEM production and cDNA libraries of the 5’ V3 gene expression and TCR kit. The remaining cell populations were also resuspended at 1000 cells/µL, but samples were not multiplexed. These samples were run on their own lane of the 10X chip for GEM production and cDNA library production using the 3’ gene expression kit. Samples were sequenced on illumina sequencer. FASTQ files generated were processed by 10X cell ranger software to generate gene counts and barcode assignment. For the multiplexed CD4+ cell samples a combination of FREEMUXLET and DEMUXLET software was used to unbiasedly identify which cells belong to each subject by single nucleotide polymorphisms in the axiom 1000 data set (Van der Auwera et al. 2013; Kang et al. 2018; Anon n.d.). The SNPs used for sample ID and the breakdown between SNPs where at least one individual in a sample had a different allele are detailed in table S1. To confirm the robustness of SNP fingerprint identified by unbiased FREEMUXLET identification, samples from a heterogenous data set were compared to both another heterogenous dataset containing the same individuals and the homogenous 3’ datasets [table S2 and 6]. These indicate that samples are consistently identified the same way in heterogeneous datasets and the SNP fingerprint matches between the 5’ and 3’ data sets.

Sample QC, integration, PCA, dimensionality reduction, clustering, marker gene, and DEG analysis were performed using the Seurat package V3 for R (Stuart et al. 2019). Single cell Gene Set Enrichment Analysis was performed using the escape package (Borcherding et al. 2021) to assign a gene set enrichment score for human tissue resident memory T cells as defined by (Kumar et al. 2017).

### Activation Induced Marker Assay

The assay protocol generally followed those provided in (Bacher & Scheffold 2013; Bacher et al. 2016). Cryopreserved BAL samples were thawed and washed once with RPMI-1640 containing 20 % FCS and 25 µg/mL DNase I. Washed samples were resuspended in 200 µL complete kool aid media plus blocking anti-CD40 antibody and seeded to 1, 96 u-bottom well per 500,000 total cells. Cells were rested for 6 hours at 37 degrees celsius in 10% CO_2_ and 100% relative humidity. After resting, cells were stimulated for 8 hours with 40 µg/mL of HDM extract or PBS vehicle. After stimulation cells were harvested, stained, and sorted for flow according to the same strategy and protocol as cryopreserved samples. During the staining step, CITE-Seq antibodies were included as defined in the subsequent table. After sorting, scRNA-seq was performed on the CD4+ T cells using the same approach as the cryopreserved samples.

### Statistics and Analytical Software

Statistical analyses and plotting was performed using Graphpad Prism (Version 9.2.0) and R (version 4.2.1).

## Data Availability

All raw sequencing data for scRNAseq and bulk blood TCRseq are deposited in GEO under BioProject ID PRJNA1477178. Raw bulk mRNA sequencing data from epithelial brushing is available in the repository GSE250241.

## Acknowledgements

We thank the UCSF Genomics CoLab for technical expertise and assistance with genomics and mass cytometry. We acknowledge the PFFC (RRID:SCR_018206) for assistance with FACS sorting, supported in part by the DRC Center Grant NIH P30 DK063720. This work was supported by the US National Institutes of Health (U19AI077439, R01HL109102, K24 HL137013), the Sandler Asthma Basic Research Center, the Hooper Foundation, the Wenner-Gren Foundations (FT2022-0006), the Swedish Heart-Lung Foundation, the Sweden-America Foundation, The Herman Krefting Foundation, Sahlgrenska Academy Starting Grant, and the Foundation Blanceflor Boncompagni Ludovisi, née Bildt. Priscila Muñoz-Sandoval is a Howard Hughes Medical Institute Gilliam Fellow.

## Supplementary Material

Figure S1 describes the hierarchical bi-variate gating strategy used to define cell types for cytof analyses. Figure S2 describes the hierarchical bi-variate gating strategy used to sort cell by flow cytometry and subsequently pool populations for downstream scRNAseq analyses. Figure S3 is a heat map displaying the phenotypic distribution of identified clones in the unstimulated scRNAseq on CD4+ T cells. Figure S4 describes the V3/V2 enrichment of *TRB* sequences in the blood of subjects broken out by corresponding unstimulated and stimulated phenotype as in Figure 5 for *TRA* sequences. Table S1 describes how many SNPs were evaluated to identify individuals via the FREEMUXLET algorithm, and how many were shared amongst individuals versus unique to an individual. Table S2 describes how many cells matched to a specific subject genotype based upon unbiased identification in the FREEMUXLET algorithm. Table S3 describes how many cells matched to an unbiasedly generated genotype based upon unbiased identification in the FREEMUXLET algorithm in two independent samples containing the same subjects. Table S4 describes detailed information on the antibodies used for cytof analysis. Table S5 describes detailed information on the antibodies used for fluorescent flow cytometry based sorting prior to scRNAseq. Table S6 describes detailed information on the CITEseq antibodies used in the AIM assay to detect protein expression via scRNA sequencing.

---

**Supplemental Figure 1 (for figure 2).**
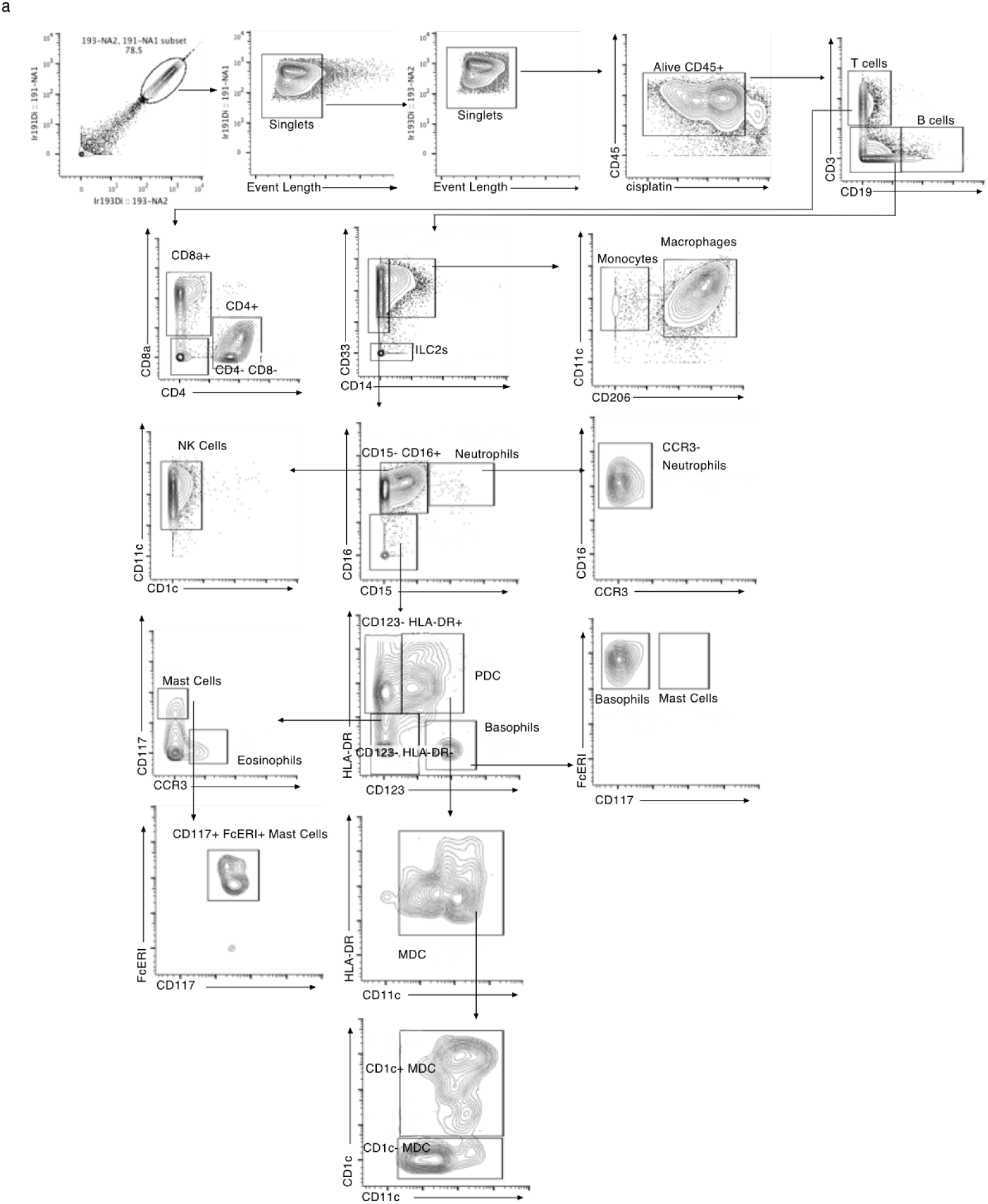
CyTOF Gating Strategy a. Gating strategy for identification of cell types in CyTOF samples. Data shown are from a single representative BAL sample.

**Supplemental Figure 2 (for Figure 4).**
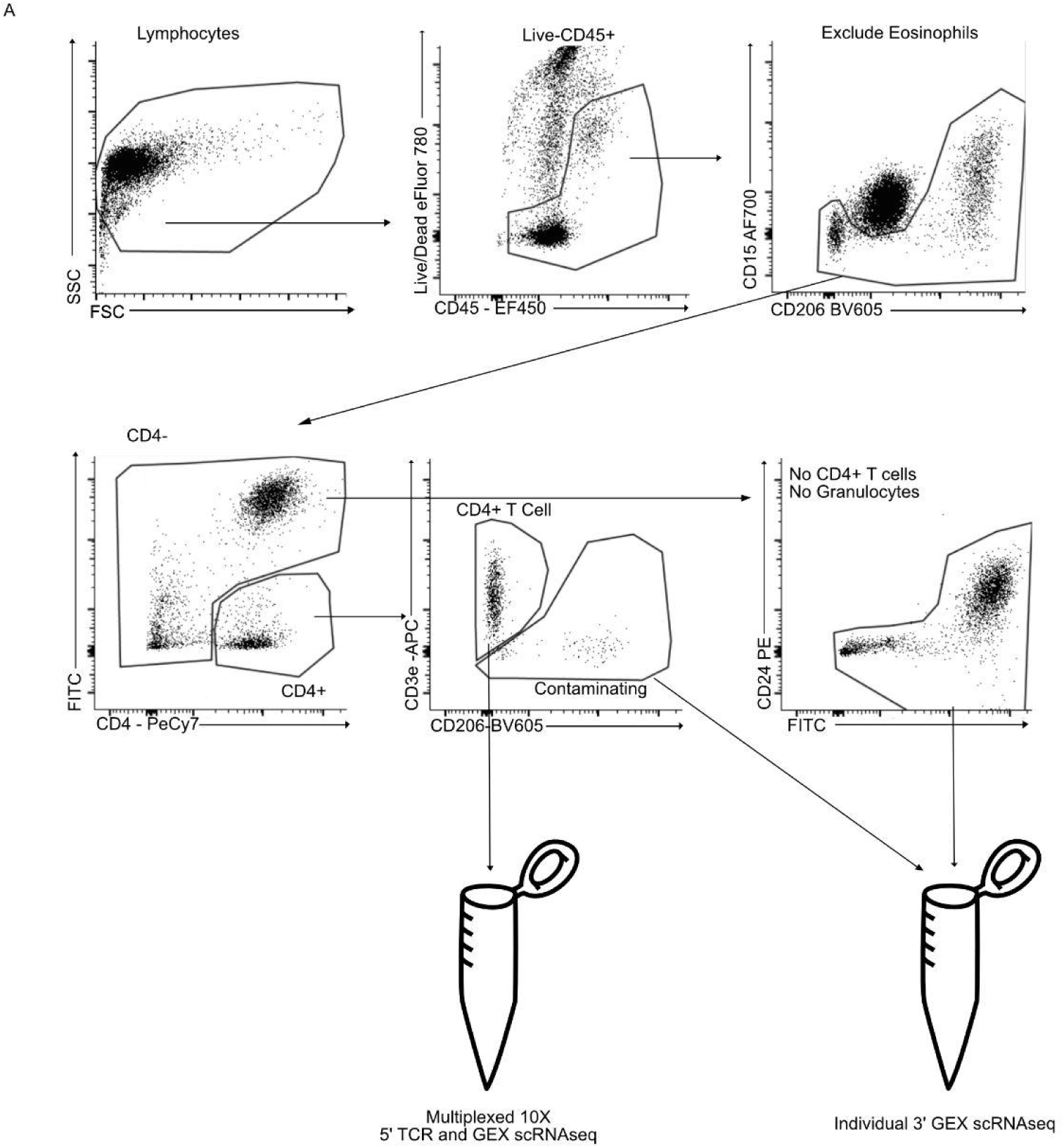
Sorting strategy for depleting granulocytes and enriching CD4+ T cells from BAL a. Gating strategy for sorting BAL from a single representative allergen challenged sample

**Supplemental Figure 3 (for Figure 5).**
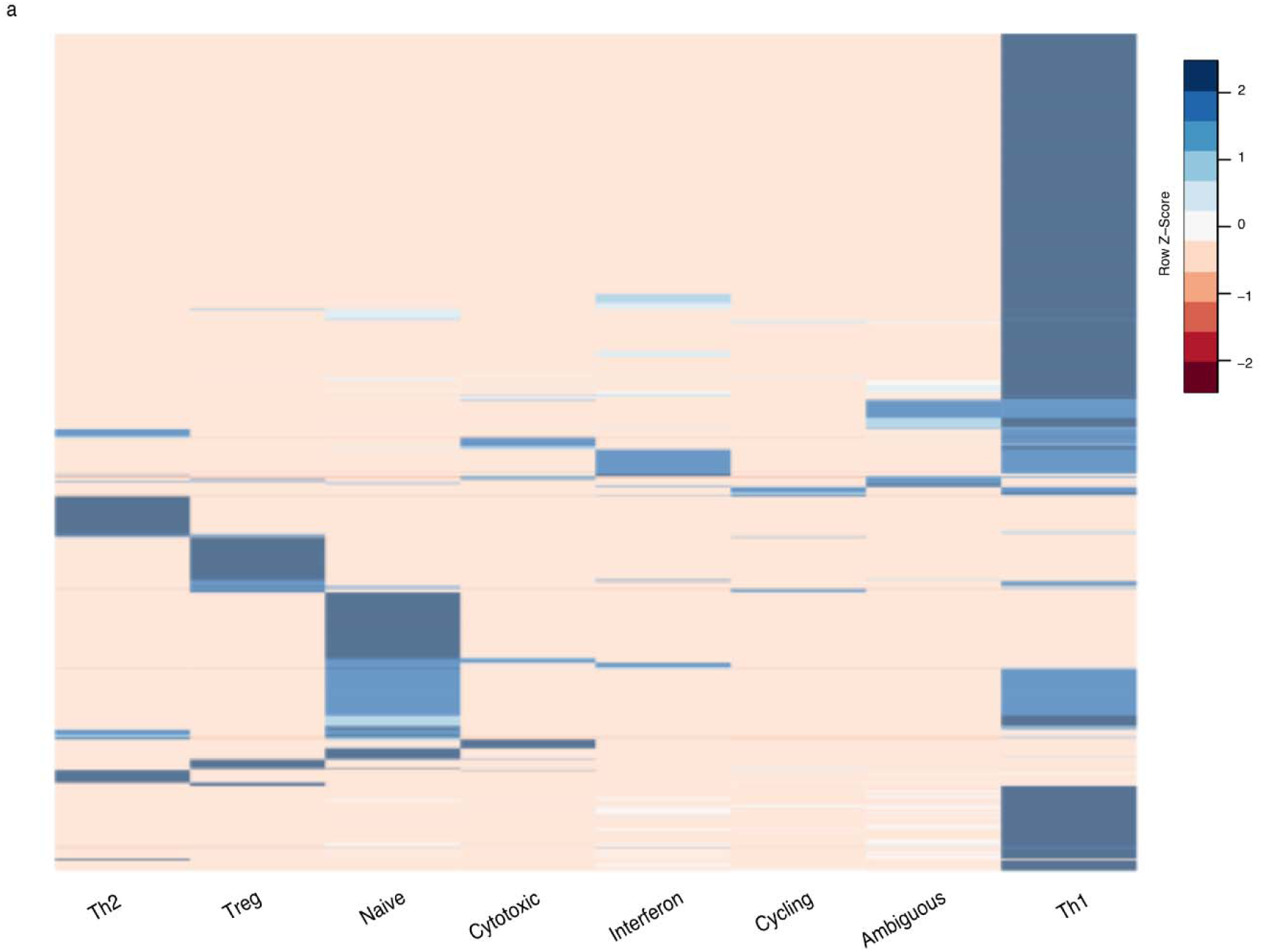
TCR clones are cluster restricted a. Heatmap indicating the frequency of phenotypes for each expanded clonotype in the directly assayed data set. Each row represents a single clonotype and the column color indicates the frequency of that cluster found in the cells of that clonotype. Rows are normalized and hierarchically clustered.

**Supplemental Figure 4 (for figure 5).**
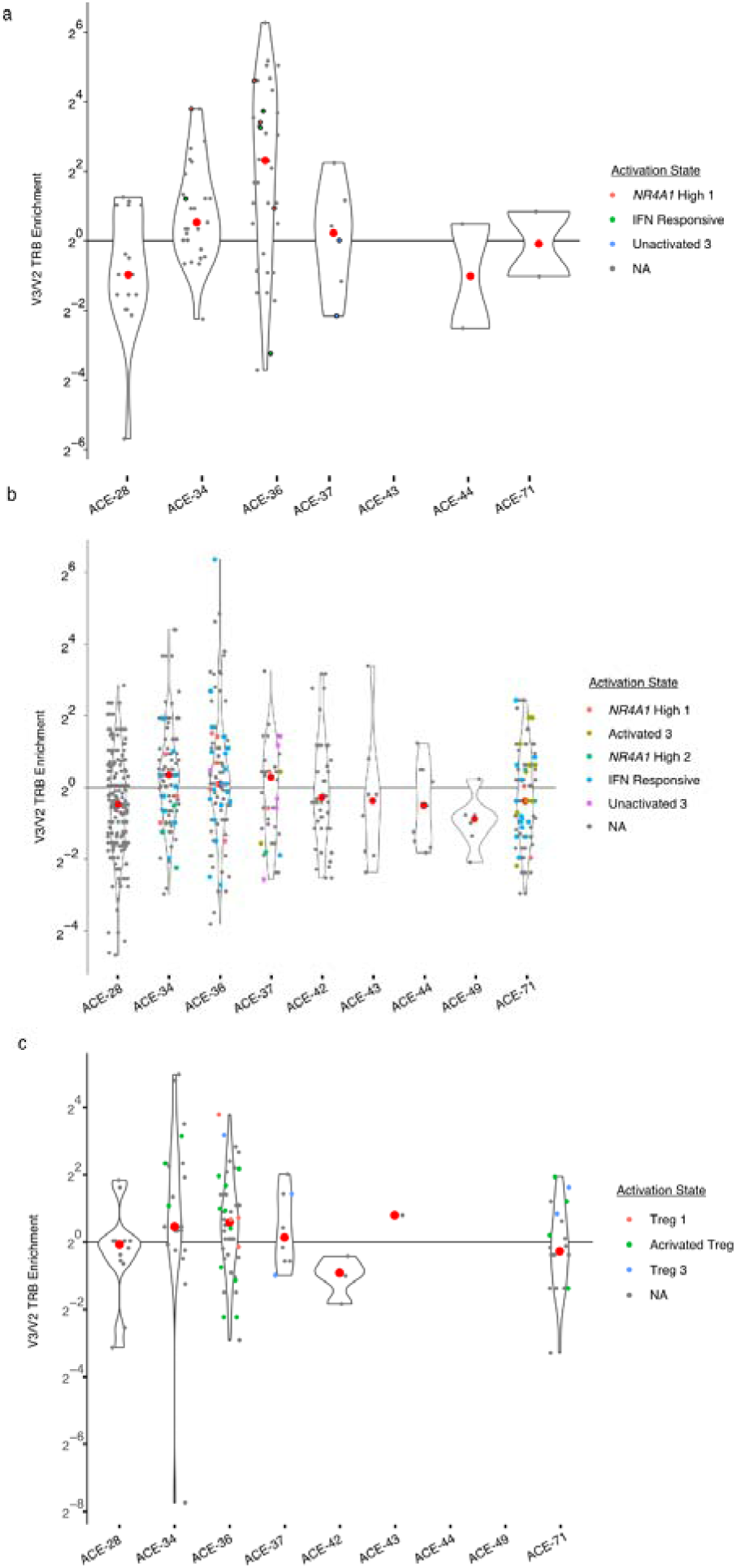
*TRB* blood enrichment mirrors *TRA* enrichment a. Enrichment of *TRB* sequences associated with clonotypes identified as Th2 clones in the directly assayed data set. Select clones that were also identified in the AIM assay are color coded by the most prominent cluster from that assay. b. Enrichment of *TRB* sequences associated with clonotypes identified as Th1 clones in the directly assayed data set. Select clones that were also identified in the AIM assay are color coded by the most prominent cluster from that assay. c. Enrichment of *TRB* sequences associated with clonotypes identified as Treg clones in the directly assayed data set. Select clones that were also identified in the AIM assay are color coded by the most prominent cluster from that assay.

**Table S1.** SNPs Used for Sample Identification.

| Total SNPs Evaluated | Unique SNPs | Shared SNPs |
| --- | --- | --- |
| 36637 | 3194 | 33443 |

**Table S2.**
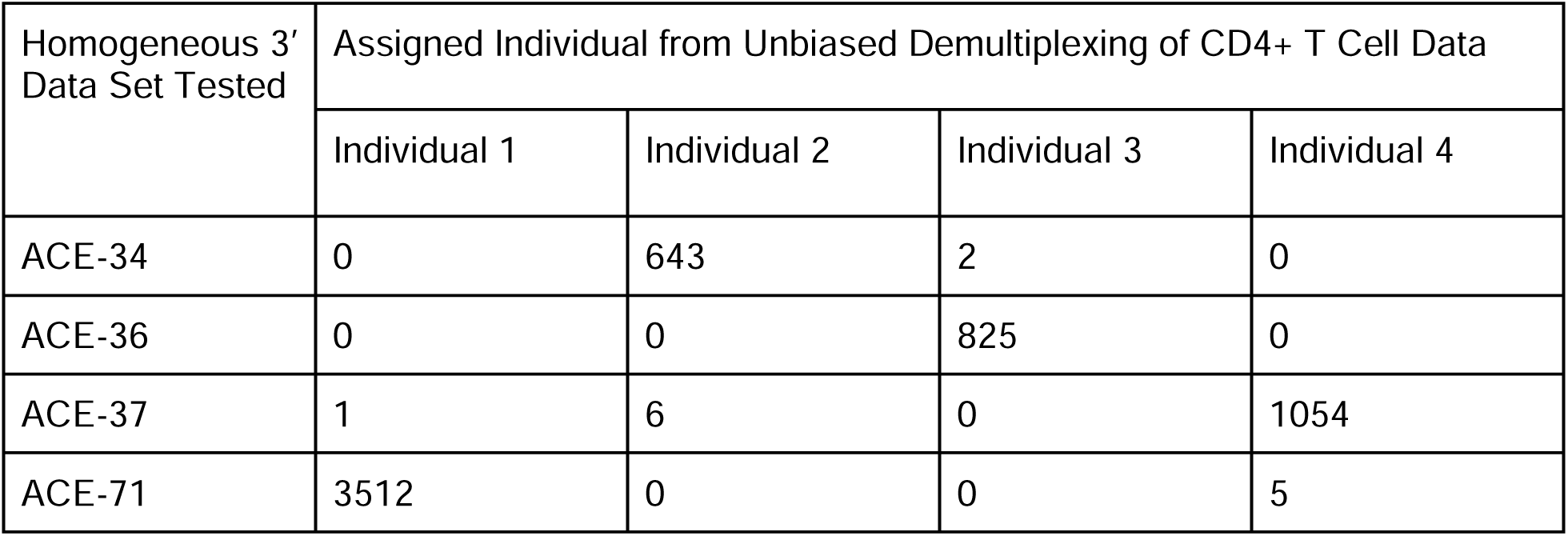
Assigning cells from homogenous data sets to individuals defined via unbiased demultiplexing of heterogeneous samples.

| Homogeneous 3' Data Set Tested | Assigned Individual from Unbiased Demultiplexing of CD4+ T Cell Data |  |  |  |
| --- | --- | --- | --- | --- |
|  | Individual 1 | Individual 2 | Individual 3 | Individual 4 |
| ACE-34 | 0 | 643 | 2 | 0 |
| ACE-36 | 0 | 0 | 825 | 0 |
| ACE-37 | 1 | 6 | 0 | 1054 |
| ACE-71 | 3512 | 0 | 0 | 5 |

**Table S3.** Cross Referencing assignments of cells from two independent heterogeneous samples containing the same individuals.

| Sample Set 2 Individual | Assigned Individual from Unbiased Demultiplexing of CD4+ T Cell Data |  |  |  |
| --- | --- | --- | --- | --- |
|  | Individual 1 | Individual 2 | Individual 3 | Individual 4 |
| Individual 1 | 2917 | 0 | 0 | 0 |
| Individual 2 | 0 | 2627 | 0 | 0 |
| Individual 3 | 1 | 0 | 2516 | 0 |
| Individual 4 | 0 | 0 | 0 | 678 |

**Table S4.**
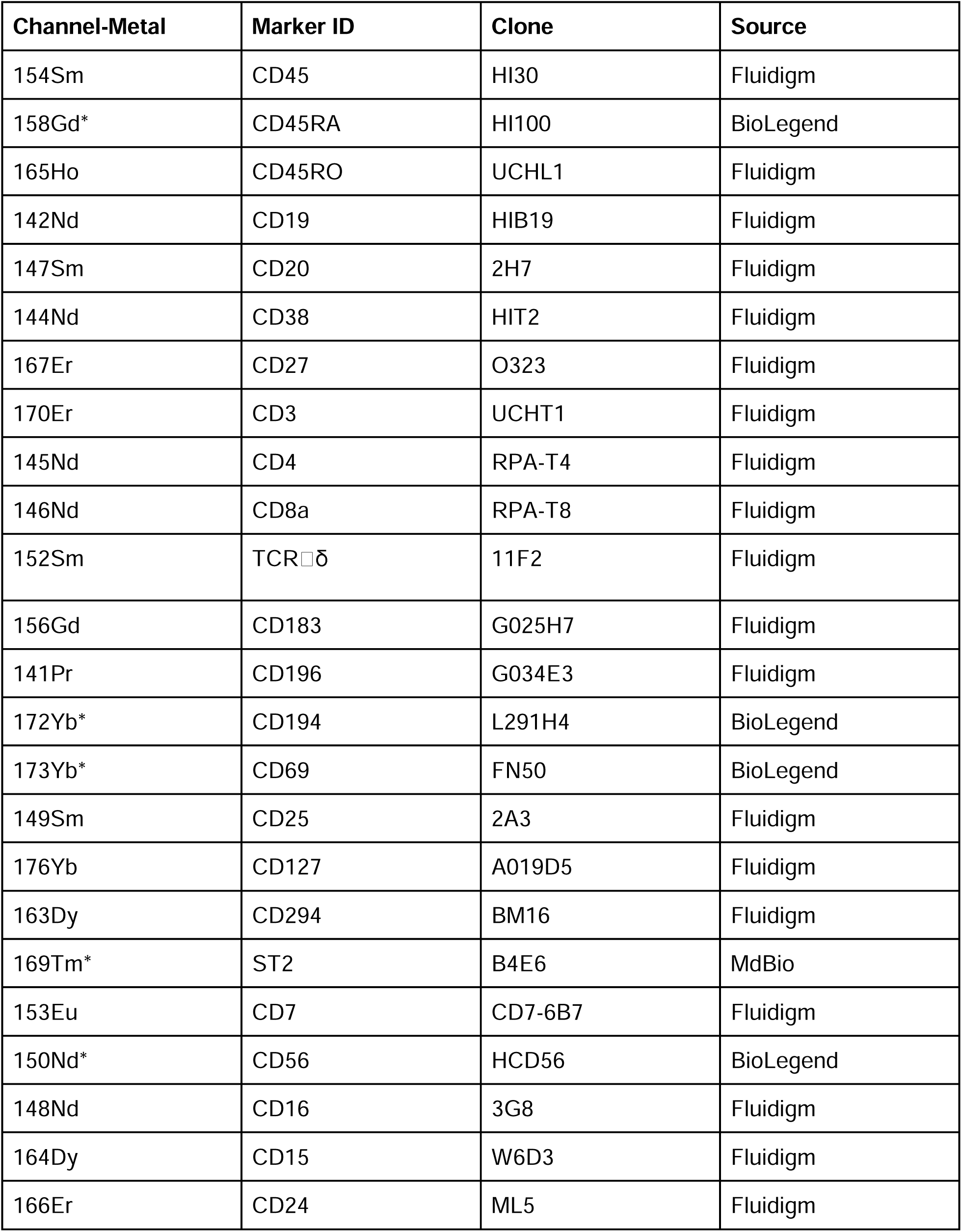
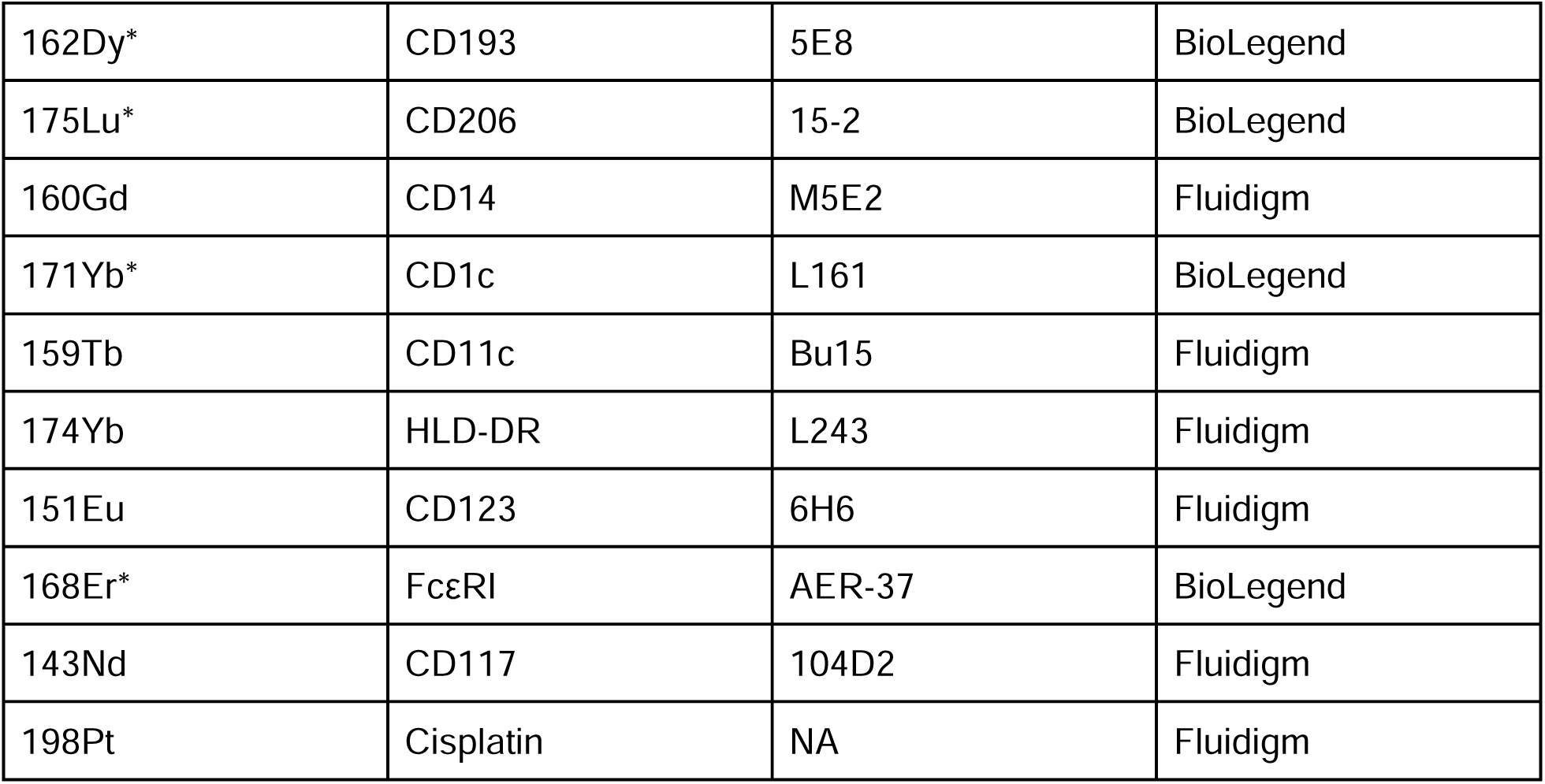
CyTOF Staining Antibodies.

| Channel-Metal | Marker ID | Clone | Source |
| --- | --- | --- | --- |
| 154Sm | CD45 | HI30 | Fluidigm |
| 158Gd* | CD45RA | HI100 | BioLegend |
| 165Ho | CD45RO | UCHL1 | Fluidigm |
| 142Nd | CD19 | HIB19 | Fluidigm |
| 147Sm | CD20 | 2H7 | Fluidigm |
| 144Nd | CD38 | HIT2 | Fluidigm |
| 167Er | CD27 | O323 | Fluidigm |
| 170Er | CD3 | UCHT1 | Fluidigm |
| 145Nd | CD4 | RPA-T4 | Fluidigm |
| 146Nd | CD8a | RPA-T8 | Fluidigm |
| 152Sm | TCR$\alpha\delta$ | 11F2 | Fluidigm |
| 156Gd | CD183 | G025H7 | Fluidigm |
| 141Pr | CD196 | G034E3 | Fluidigm |
| 172Yb* | CD194 | L291H4 | BioLegend |
| 173Yb* | CD69 | FN50 | BioLegend |
| 149Sm | CD25 | 2A3 | Fluidigm |
| 176Yb | CD127 | A019D5 | Fluidigm |
| 163Dy | CD294 | BM16 | Fluidigm |
| 169Tm* | ST2 | B4E6 | MdBio |
| 153Eu | CD7 | CD7-6B7 | Fluidigm |
| 150Nd* | CD56 | HCD56 | BioLegend |
| 148Nd | CD16 | 3G8 | Fluidigm |
| 164Dy | CD15 | W6D3 | Fluidigm |
| 166Er | CD24 | ML5 | Fluidigm |
| 162Dy* | CD193 | 5E8 | BioLegend |
| 175Lu* | CD206 | 15-2 | BioLegend |
| 160Gd | CD14 | M5E2 | Fluidigm |
| 171Yb* | CD1c | L161 | BioLegend |
| 159Tb | CD11c | Bu15 | Fluidigm |
| 174Yb | HLD-DR | L243 | Fluidigm |
| 151Eu | CD123 | 6H6 | Fluidigm |
| 168Er* | FcεRI | AER-37 | BioLegend |
| 143Nd | CD117 | 104D2 | Fluidigm |
| 198Pt | Cisplatin | NA | Fluidigm |

**Table S5.** FACS Staining Antibodies.

| Reagent | Source | Detail | Catalog # |
| --- | --- | --- | --- |
| CD4 - PE-Cy7 | Biolegend | Clone OKT4 | 317413 |
| CD45 - ef450 | Invitrogen | 2D1 | 48-9459-42 |
| CD24 PE | BD Biosciences | ML5 | 560991 |
| CD206 BV605 | BD Biosciences | 19.2 | 740417 |
| CD3 APC | Biolegend | UCHT1 | 300412 |
| CD15 AF700 | Biolegend | HI98 | 301919 |
| CD20 FITC | Biolegend | 2H7 | 302303 |
| CD19 FITC | Biolegend | HIB19 | 302206 |
| CD16 FITC | Biolegend | 3G8 | 302005 |
| CD14 FITC | Biolegend | HCD14 | 325603 |
| CD56 FITC | Biolegend | MEM-188 | 304603 |

**Table S6.**
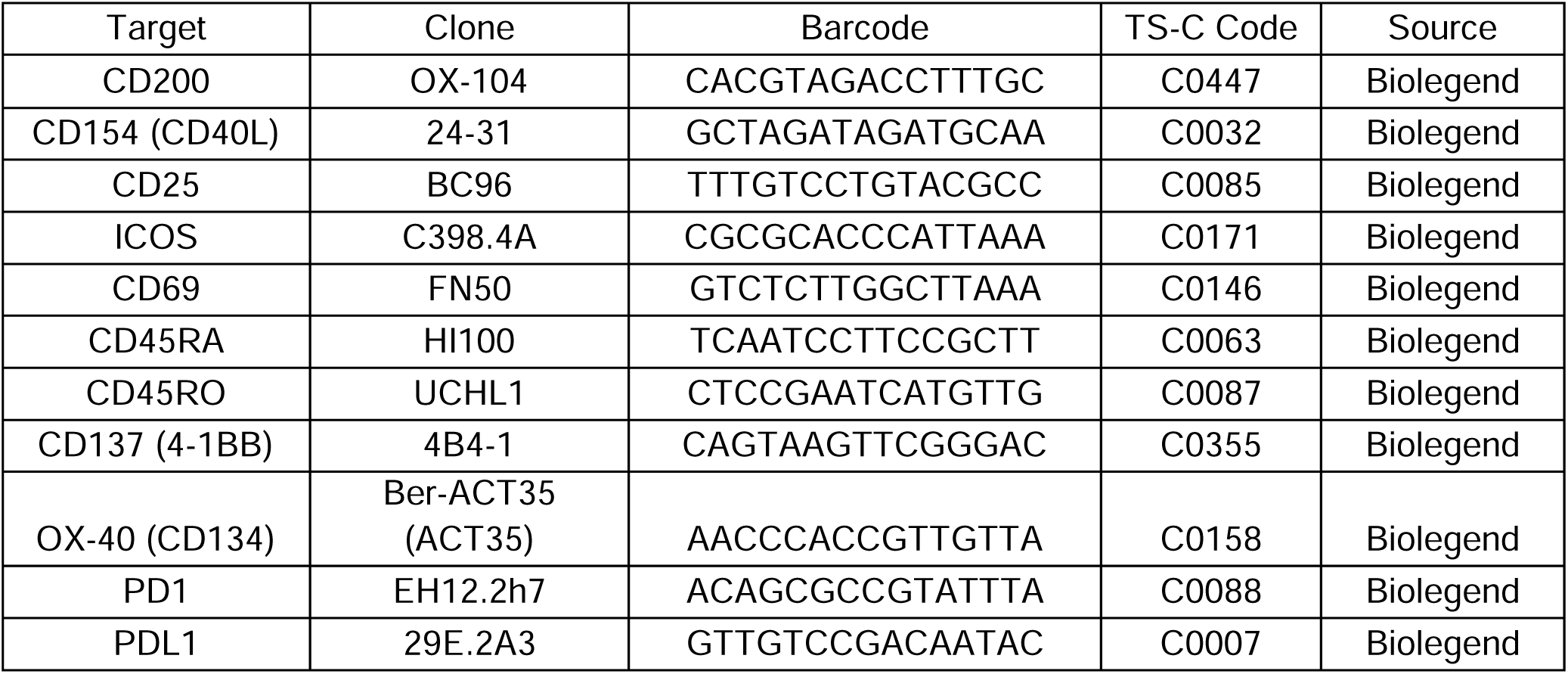
CITE-Seq Staining Antibodies.

## References

Alladina, J. et al., 2023. A human model of asthma exacerbation reveals transcriptional programs and cell circuits specific to allergic asthma. Science immunology, 8(83), p.eabq6352.

Anon, popscle: A suite of population scale analysis tools for single-cell genomics data including implementation of Demuxlet / Freemuxlet methods and auxilary tools, Github. Available at: https://github.com/statgen/popscle [Accessed May 14, 2023].

Bacher, P. et al., 2016. Regulatory T Cell Specificity Directs Tolerance versus Allergy against Aeroantigens in Humans. Cell, 167(4), pp.1067–1078.e16.

Bacher, P. & Scheffold, A., 2013. Flow-cytometric analysis of rare antigen-specific T cells. Cytometry. Part A: the journal of the International Society for Analytical Cytology, 83(8), pp.692–701.

Bhakta, N.R. et al., 2013. A qPCR-based metric of Th2 airway inflammation in asthma. Clinical and translational allergy, 3(1), p.24.

Borcherding, N. et al., 2021. Mapping the immune environment in clear cell renal carcinoma by single-cell genomics. Communications Biology, 4(1), p.122.

Castro, M. et al., 2018. Dupilumab Efficacy and Safety in Moderate-to-Severe Uncontrolled Asthma. The New England journal of medicine, 378(26), pp.2486–2496.

Cho, J.L. et al., 2016. Allergic asthma is distinguished by sensitivity of allergen-specific CD4+ T cells and airway structural cells to type 2 inflammation. Science translational medicine, 8(359), p.359ra132.

Collin, M. & Bigley, V., 2018. Human dendritic cell subsets: an update. Immunology, 154(1), pp.3–20.

Duchesne, M., Okoye, I. & Lacy, P., 2022. Epithelial cell alarmin cytokines: Frontline mediators of the asthma inflammatory response. Frontiers in immunology, 13, p.975914.

Dunican, E.M. et al., 2018. Mucus plugs in patients with asthma linked to eosinophilia and airflow obstruction. The Journal of clinical investigation, 128(3), pp.997–1009.

Fitzpatrick, A.M. et al., 2011. Heterogeneity of severe asthma in childhood: confirmation by cluster analysis of children in the National Institutes of Health/National Heart, Lung, and Blood Institute Severe Asthma Research Program. The Journal of allergy and clinical immunology, 127(2), pp.382–389.e1–13.

Gauvreau, G.M. et al., 2014. Effects of an anti-TSLP antibody on allergen-induced asthmatic responses. The New England journal of medicine, 370(22), pp.2102–2110.

GBD Chronic Respiratory Disease Collaborators, 2020. Prevalence and attributable health burden of chronic respiratory diseases, 1990-2017: a systematic analysis for the Global Burden of Disease Study 2017. The Lancet. Respiratory medicine, 8(6), pp.585–596.

Haldar, P. et al., 2008. Cluster analysis and clinical asthma phenotypes. American journal of respiratory and critical care medicine, 178(3), pp.218–224.

Hansen, S. et al., 2024. Clinical response and remission in patients with severe asthma treated with biologic therapies. Chest, 165(2), pp.253–266.

Hartoularos, G.C. et al., 2023. Reference-free multiplexed single-cell sequencing identifies genetic modifiers of the human immune response. bioRxiv, p.2023.05. 29.542756. Available at: https://www.biorxiv.org/content/10.1101/2023.05.29.542756.abstract.

Herzog, S. et al., 2022. Myeloid CD169/Siglec1: An immunoregulatory biomarker in viral disease. Frontiers of medicine, 9, p.979373.

Hirai, H. et al., 2001. Prostaglandin D2 selectively induces chemotaxis in T helper type 2 cells, eosinophils, and basophils via seven-transmembrane receptor CRTH2. The Journal of experimental medicine, 193(2), pp.255–261.

Jheng, M.-J. et al., 2023. Chemokine Receptor CCR8 and CCR8-Bearing Cells Are Involved in Regulation of Type 2 Immunity in Response to Airborne Allergens. The Journal of allergy and clinical immunology, 151(2), p.AB219.

Kang, H.M. et al., 2018. Multiplexed droplet single-cell RNA-sequencing using natural genetic variation. Nature biotechnology, 36(1), pp.89–94.

Koh, K.D. et al., 2023. Genomic characterization and therapeutic utilization of IL-13-responsive sequences in asthma. Cell genomics, 3(1), p.100229.

Kumar, B.V. et al., 2017. Human Tissue-Resident Memory T Cells Are Defined by Core Transcriptional and Functional Signatures in Lymphoid and Mucosal Sites. Cell reports, 20(12), pp.2921–2934.

Lambrecht, B.N., Hammad, H. & Fahy, J.V., 2019. The Cytokines of Asthma. Immunity, 50(4), pp.975–991.

Lei, H. et al., 2021. Single-cell RNA-Seq revealed profound immune alteration in the peripheral blood of patients with bacterial infection. International journal of infectious diseases: IJID: official publication of the International Society for Infectious Diseases, 103, pp.527–535.

Liang, X. et al., 2019. Macrophage FABP4 is required for neutrophil recruitment and bacterial clearance in Pseudomonas aeruginosa pneumonia. FASEB journal: official publication of the Federation of American Societies for Experimental Biology, 33(3), pp.3562–3574.

Lin, J. et al., 2018. Symbicort® Maintenance and Reliever Therapy (SMART) and the evolution of asthma management within the GINA guidelines. Expert review of respiratory medicine, 12(3), pp.191–202.

Lloyd, C.M. & Hessel, E.M., 2010. Functions of T cells in asthma: more than just T(H)2 cells. Nature reviews. Immunology, 10(12), pp.838–848.

Martin, J.C. et al., 2019. Single-Cell Analysis of Crohn’s Disease Lesions Identifies a Pathogenic Cellular Module Associated with Resistance to Anti-TNF Therapy. Cell, 178(6), pp.1493–1508.e20.

Melgert, B.N. et al., 2007. Effects of 4 months of smoking in mice with ovalbumin-induced airway inflammation. Clinical and experimental allergy: journal of the British Society for Allergy and Clinical Immunology, 37(12), pp.1798–1808.

Menzies-Gow, A. et al., 2003. Anti-IL-5 (mepolizumab) therapy induces bone marrow eosinophil maturational arrest and decreases eosinophil progenitors in the bronchial mucosa of atopic asthmatics. The Journal of allergy and clinical immunology, 111(4), pp.714–719.

Moore, W.C. et al., 2010. Identification of asthma phenotypes using cluster analysis in the Severe Asthma Research Program. American journal of respiratory and critical care medicine, 181(4), pp.315–323.

Moran, A. & Pavord, I.D., 2020. Anti-IL-4/IL-13 for the treatment of asthma: the story so far. Expert opinion on biological therapy, 20(3), pp.283–294.

Nassar, A.F., Wisnewski, A.V. & Raddassi, K., 2016. Mass cytometry moving forward in support of clinical research: advantages and considerations. Bioanalysis, 8(4), pp.255–257.

Padem, N. & Saltoun, C., 2019. Classification of asthma. Allergy and asthma proceedings: the official journal of regional and state allergy societies, 40(6), pp.385–388.

Pilette, C. et al., 2004. CCR4 ligands are up-regulated in the airways of atopic asthmatics after segmental allergen challenge. The European respiratory journal: official journal of the European Society for Clinical Respiratory Physiology, 23(6), pp.876–884.

Rahimi, R.A. et al., 2026. Distinct phenotypes and repertoires of bronchoalveolar and airway mucosal T cells in health and allergic asthma. Mucosal Immunology. Available at: 10.1016/j.mucimm.2026.01.001 [Accessed April 6, 2026].

Saggau, C., Scheffold, A. & Bacher, P., 2021. Flow cytometric characterization of human antigen-reactive T-helper cells. *Methods in molecular biology (Clifton*, N.J.), 2285, pp.141– 152.

Saradna, A. et al., 2018. Macrophage polarization and allergic asthma. Translational research: the journal of laboratory and clinical medicine, 191, pp.1–14.

Seumois, G. et al., 2020. Single-cell transcriptomic analysis of allergen-specific T cells in allergy and asthma. Science Immunology, 5(48), p.eaba6087.

Strazza, M. & Mor, A., 2017. Consider the Chemokines: a Review of the Interplay Between Chemokines and T Cell Subset Function. Discovery medicine, 24(130), pp.31–39.

Stuart, T. et al., 2019. Comprehensive Integration of Single-Cell Data. Cell, 177(7), pp.1888– 1902.e21.

Tang, M. et al., 2022. Mucus Plugs Persist in Asthma, and Changes in Mucus Plugs Associate with Changes in Airflow over Time. American journal of respiratory and critical care medicine, 205(9), pp.1036–1045.

Van der Auwera, G.A., et al., 2013. From FastQ data to high confidence variant calls: the Genome Analysis Toolkit best practices pipeline. Current protocols in bioinformatics / editoral board, Andreas D. Baxevanis … [et al.], 43(1110), pp.11.10.1–11.10.33.

Wenzel, S. et al., 2016. Dupilumab efficacy and safety in adults with uncontrolled persistent asthma despite use of medium-to-high-dose inhaled corticosteroids plus a long-acting β2 agonist: a randomised double-blind placebo-controlled pivotal phase 2b dose-ranging trial. The Lancet, 388(10039), pp.31–44.

Woodruff, P.G. et al., 2009. T-helper type 2-driven inflammation defines major subphenotypes of asthma. American journal of respiratory and critical care medicine, 180(5), pp.388–395.

